# Hyaluronan and skin elasticity: Are subterranean mammals special?

**DOI:** 10.64898/2026.02.11.705254

**Authors:** Kai R Caspar, Delphine del Marmol, Lisa Gerdes, Anne Zockoll, Stefanie Schuelpen, Sabine Begall

## Abstract

It has been hypothesized that subterranean mammals have evolved increased skin elasticity to reduce friction when moving through their underground tunnel systems. This trait is commonly believed to be mediated by greatly elongated hyaluronan (HA) polymers in the dermal extracellular matrix, which have been reported from different distantly related burrowing mammals. However, replicating these findings has proven difficult, and a mechanism by which HA polymer size could modify skin elasticity has not been proposed. In fact, experimental data on skin biomechanics in burrowing mammals are currently unavailable. Here, we quantify the molecular mass of HA polymers extracted from the tissues of Ansell’s mole-rat (*Fukomys anselli*), a burrowing rodent yet unstudied in that respect, and investigate skin biomechanics in subterranean and epigeic (aboveground foraging) small mammal species by means of *in vivo*-cutometry.

We did not recover abundant extremely elongated HA polymers in Ansell’s mole-rat, conflicting with published findings in congeneric species and the naked mole-rat. Polymer length was found to be increased compared to guinea pigs across tissues, though. Our data on skin biomechanics indicate that subterranean mammal skin is not consistently more elastic than that of epigeic forms. Interestingly, the skin of the naked mole-rat was characterized by very high stiffness. Uniaxial tensile tests demonstrated that it also exhibits exceptional tensile strength. Hence, we challenge the idea that hyaluronan or a subterranean ecology notably influences skin elasticity in small mammals and suggest that previous studies may have confused elasticity with skin looseness, which constitutes a different phenomenon.

## Introduction

Biomechanical adaptations of mammalian skin have so far received little attention in comparative research. Despite that, it is known that the integument of various mammals shows peculiar structural characteristics related to their ecology (Jarman, 1989; Mo et al., 2020; Seifert et al., 2012; Shadwick et al., 1992; Swartz et al., 1996). In light of this, it would not be surprising if the integument of subterranean mammals would also exhibit specialized traits facilitating locomotion in their tunnel systems and supporting homeostasis underground. Indeed, it has been noted that the fur of underground-living mammals exhibits structural peculiarities relating to increased pliability as well as resilience (Chernova & Zherebtsova, 2022; Klauer et al, 1997; Krmpotic et al., 2024). Depending on digging-mode, various species also display local thickening of the skin in body regions that experience high friction during burrowing (Klauer et al., 1997; Sokolov, 1982). Comparative molecular approaches have shown that genes coding for foot pad keratin and fibrillary proteins of the epidermal basal lamina are notably altered across subterranean lineages (Davies et al., 2018; Partha et al., 2017; Zhao et al., 2023). Otherwise, the skin of burrowing mammals typically exhibits weakly developed subcutaneous connective tissue that lacks notable fat deposits (e.g., Daly & Buffenstein, 1998). This trait supports effective heat dissipation (Pleštilová et al., 2020) and enables skin looseness (Menton et al., 1978). The latter term refers to the skin being markedly mobile relative to the underlying musculature (Tucker, 1981). Loose skin reduces abrasion of the integument during digging and evolved convergently in various groups of burrowing tetrapods (Gans, 1978; Summers & O’Reilly, 1997). However, since it is also prominently present in small mammals in general (Kawamata et al., 2003) it might well be a plesiomorphic trait that is simply retained in subterranean mammal lineages.

Here, we focus on another potential skin adaptation in underground-dwelling mammals: unusually long hyaluronan (HA) polymers in the dermis. In a highly influential paper, Tian et al. (2013) speculated that subterranean rodents evolved such conspicuously elongated HA polymers to enable increased skin elasticity. This in turn was proposed to reduce friction and improve agility in narrow underground tunnels. HA is an important mucopolysaccharide component of the vertebrate extracellular matrix. It is found across all tissue types, but it is particularly abundant in the dermis (Armstrong & Bell, 2002; Cowman et al., 2015). It is an evenly negatively-charged, non-branching polymer that consists of non-modified disaccharide units, thus displaying a uniform chemical make-up across tissues. However, depending on tissue type and species, the length and concentration of HA polymers, quantified as differences in molecular mass, can vary dramatically (Cowman, et al., 2015). Such molecular mass disparity is not trivial, because it affects the hydration and viscosity of the extracellular matrix. Precisely, longer polymers are assumed to increase the viscosity of the extracellular matrix, binding interstitial water more effectively (Cowman et al., 2015; Tian et al., 2013).

Initially, Tian et al. (2013) reported dermal HA molecules of exceptional molecular mass (6-12 MDa) in naked mole-rats (*Heterocephalus glaber* RÜPPEL, 1842-NMR) and Middle East blind mole-rats (*Nannospalax ehrenbergi* NEHRING, 1898, previously classified as *Spalax galili* NEVO ET AL., 2001). In other small mammals such polymers were found to be, for the most part, substantially smaller (0.5-7 MDa) than that (Armstrong & Bell, 2002; Reed et al., 1988; Tian et al., 2013). More recently, Zhao et al. (2023) reported that further subterranean mammals, including rodents as well as talpid moles produce HA of exceptionally high molecular mass in their tissues, especially the skin (Zhao et al., 2023). Since the species investigated in this study represent several lineages that independently adapted to underground life, the authors echo the hypothesis that subterranean mammals evolved elongated polymers convergently “to provide elastic skin that facilitates squeezing into underground tunnels” (Zhao et al., 2023). Yet, interestingly, all species screened by Zhao et al. (2023) showed a systemic increase in HA polymer length across all studied tissues and not just the skin.

The skin-elasticity hypothesis has remained untested but still found its way into numerous articles on NMRs specifically (Kulaberoglu et al., 2019; Lagunas-Rangel, 2024; Lagunas-Rangel & Chávez-Valencia, 2017; Menon et al., 2019). Unfortunately, it remains unclear what exactly the aforementioned studies are referring to when applying the term *skin elasticity*. Three different aspects of skin biomechanics could potentially be concerned here (Fig. 1): Firstly, elastic pliability of the skin acting against vertical deformation such as suction or indentation. Secondly, tensile elasticity relevant to horizontal tissue straining. These two aspects need to be cautiously differentiated and are influenced by different structural determinants (Silver et al., 2003). Thirdly, skin looseness, the degree to which the skin adheres to the underlying musculature, might have been confused with skin elasticity.

**Figure 1:**
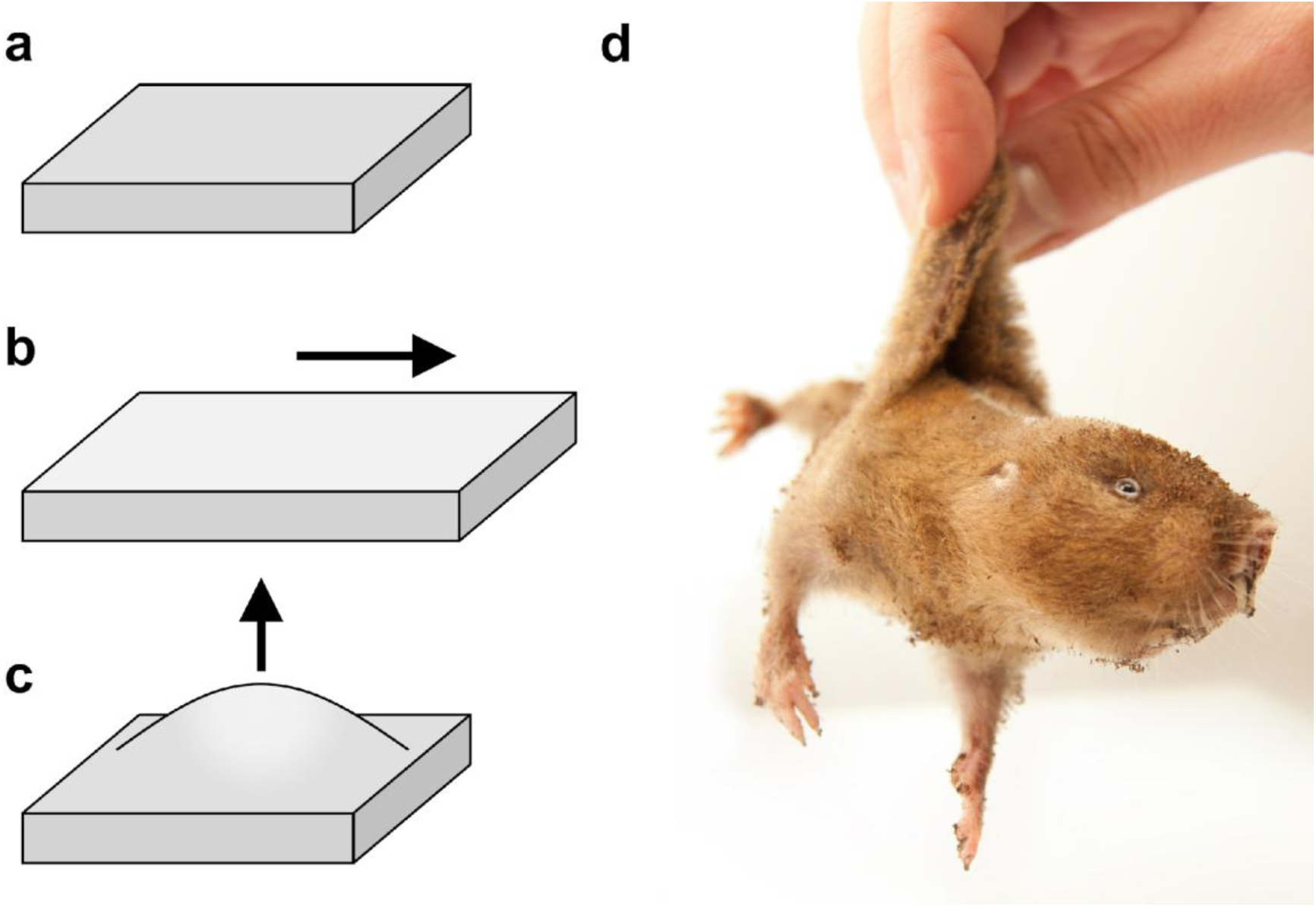
Types of skin conformation changes relevant to the aims of this study. **a**: Schematic of skin at rest in the absence of forces changing its conformation. **b**: Schematic of skin under horizontal strain (typically studied by means of uniaxial tensile straining tests). **c:** Schematic of skin experiencing vertical deformation, as induced by suction (studied here by aid of a Cutometer®). **d**: Photograph illustrating pronounced skin looseness in an Ansell’s mole-rat (*Fukomys anselli*). The skin is easily movable relative to the underlying musculature due to limited connective tissue anchoring the dermis. However, actual deformation of the skin tissue is minimal. Photo credit: Sarah M. Wilms, used with permission.

Unfortunately, experimental comparative studies addressing to what extent skin biomechanics might be altered by HA polymers of varying length and whether they actually differ between subterranean and epigeic groups (i.e. those that forage above ground) are lacking. Furthermore, important caveats need to be kept in mind when hypothesizing about dermal HA functions in burrowing mammals based on current evidence: The finding of exceptionally long HA polymers in the skin of the NMR could so far not be reliably replicated by groups independent from the authors of the Tian et al. (2013) paper (Braude et al., 2021; del Marmol et al., 2021) (but see Zhao et al. (2019); note that we are not aware of any independent attempts to replicate findings in *Nannospalax*). All studies reporting such elongated polymers so far relied on pulse-field gel electrophoresis and Alcian blue staining to identify them (Tian et al., 2013; Zhao et al., 2019; Zhao et al., 2023). However, Alcian blue does not specifically stain HA, making it unreliable to detect it and the suitability of pulse-field gel electrophoresis as a method to determine HA polymer size has been questioned (see detailed discussion by del Marmol et al. (2021)). Evidence for extremely elongated HA polymers in subterranean mammals is thus not solid. On the other hand, to the best of our knowledge, no mechanism for how HA might determine skin elasticity at the molecular level has been proposed. Studies that have tested whether HA influences the tensile properties of the dermis and other connective tissues in mammals report only negligible effects (Kronick & Sacks, 1994; Oxlund & Andreassen, 1980; Partington & Wood, 1963). This strongly suggests that increases in dermal HA polymer size do not enhance skin elasticity during horizontal straining. We are not aware of studies that tested for an effect of HA polymer size on elastic pliability of dermal tissue. In light of these issues, the hyaluronan skin elasticity-hypothesis appears not well-supported.

Here, we attempt to experimentally test the notion that skin elasticity correlates with ecology by determining the viscoelastic properties of skin from subterranean and epigeic rodent species *in vivo*. For this, we used a suction-probe equipped with optical sensors for measuring how the tissue rebounds when subjected to minor vertically applied strains. This methodology is commonly applied in research on cosmetic interventions and reconstructive surgery (Müller et al., 2018; Stroumza et al., 2015).

Additionally, we analyzed the distribution and molecular mass of HA in tissues of the Ansell’s mole-rat (*Fukomys anselli* BURDA ET AL., 1999 – AMR) to characterize an underground-dwelling rodent so far unstudied in this respect (note that it has recently been suggested that *F. anselli* should be synonymized with *F. micklemi* CHUBB, 1909 – Šumbera et al., 2026). Like its more popular naked relative, the AMR is part of the fully subterranean African mole-rat radiation (Bathyergidae) but while the NMR represents the most basal genus of the group, the AMR occupies a derived position in the bathyergid family tree (Ingram et al., 2004). Both species are long-lived, reaching lifespans of ≥ 20 years in captivity and showing notable cancer resistance (Begall et al., 2021; Braude et al., 2021). Zhao et al. (2023) have recently reported exceptionally long HA polymers in a congener of the AMR, the Damaraland mole-rat (*Fukomys damarensis* OGILBY, 1838) of the Kalahari region. Their data suggest that Damaraland mole-rats express even longer HA polymers than NMR and show extreme relative mRNA expression levels for enzymes synthesizing (HAS2: 15 times higher than in NMR, 45 times higher than in guinea pigs) and degrading HA of high molecular mass (HYAL2: 3 times higher than in NMR, 6 times higher than in guinea pigs). Given these results, similar patterns would be expected in the closely-related AMR.

## Materials and Methods

All statistics were performed in R (R Core Team, 2020). Linear mixed effect models (LMM) were computed by aid of the *lme4* package (Bates et al., 2015).

The supplementary information for this article contains a main file with three parts (part I: hyaluronan polymers (Figs. S1 and S2); part II: elastic pliability (Tables S1 and S2); part III: uniaxial tensile testing of rodent skin (Figs. S3 and S4, Tables S3 and S4)) and a table compiling data on skin pliability used for statistical analyses (Table S5).

### Hyaluronan polymer size and distribution in Ansell’s mole-rat tissues

We sampled four young adult Ansell’s mole-rats (age: 12 to 27 months; one female, three males) derived from the breeding stock of the Department of General Zoology at the University of Duisburg-Essen, Germany. We took samples of skin, skeletal muscle, and kidney immediately after sacrificing the animals (following del Marmol et al.,2021). Tissue samples used for histology were fixed in 4 % formaldehyde solution, while those used to determine HA content and molecular mass were frozen in liquid nitrogen. Analytical and histological procedures as well as reagents were identical to the ones described for NMRs in del Marmol et al. (2021) and are described in detail there.

Briefly, histological sections were prepared following a standard protocol. They were subsequently incubated in a buffer solution containing biotinylated hyaluronan binding proteins (HABP). HABP bind to hyaluronan exclusively, allowing a high signal specificity. Following incubation, HABP were detected via horseradish peroxidase-conjugated streptavidin and 3,3′-diaminobenzidine. Counterstaining was achieved with hematoxylin, and DPX was used to mount sections.

Molecular mass of HA from AMR tissue extracts (skin: *n* = 3; skeletal muscle: *n* = 4; kidney: *n* = 3) was determined by Fast Protein Liquid Chromatography-Size Exclusion Chromatography (FPLC-SEC) with a Sephacryl S-1000 (GE Healthcare Europe GmbH, Diegem, Belgium) glass column (length 26 cm, diameter 0.6 cm). HA samples with a molecular mass of 2500 kDa and 150 kDa (SelectHA; Hyalose LLC, San Diego, CA, USA), respectively, were used as standards (Figure S1). Samples were eluted at 70 µl/min with 0.15 M sodium acetate, 0.1% 3-[(3-cholamidopropyl)dimethylammonio]-1-propanesulfonate hydrate (CHAPS; Sigma-Aldrich, St. Louis, MO, USA). For each tissue extract, 30 fractions, 200 µl each, were collected. For each sample fraction, the HA concentration was measured via solid phase sandwich ELISA using Hyaluronan DuoSet kits (R&Dsystems, Abingdon, United Kingdom) following the manufacturer’s instructions and calculated as a percentage of the total HA recovered.

We specifically compare our data on AMR with published results on other hystricomorph rodents, the guinea pig (*Cavia porcellus* LINNAEUS, 1758) and the NMR by del Marmol et al. (2021) who followed the same experimental protocols. However, there was not enough usable material to separate kidney medulla and cortex for hyaluronan-related measurements in the AMR, in contrast to the two species studied previously (Figure S2). To avoid biases, we refrained from statistical comparisons of HA content in the tissues of these species because the measurements of HA levels were conducted in separate ELISA runs, the results of which may not be comparable. Limited sample sizes further prevented us from drawing meaningful statistical comparisons for HA molecular weight distributions at the interspecific level. We therefore compared HA concentrations and polymer size distributions qualitatively rather than quantitatively.

### Elastic pliability of rodent skin (*in vivo* Cutometer® tests)

#### Subjects and housing

We used three epigeic (laboratory house mouse (*Mus musculus* LINNAEUS, 1758 - C57BL/6, B6); laboratory Norway rat (*Rattus norvegicus* BERKENHOUT, 1769 - Lewis strain (LEW/Crl), LR), and domestic guinea pig (multi-colored shorthaired; GP)) and four subterranean rodent species (Ansell’s mole-rat (AMR), giant mole-rat (*Fukomys mechowii* PETERS, 1881; GMR), naked mole-rat (NMR), coruro (*Spalacopus cyanus* MOLINA, 1782; SC)) for *in vivo* vertical skin straining tests. The coruro is a social subterranean rodent, closely related to degus and endemic to Chile, which exhibits less extreme specializations related to the underground habitat than African mole-rats do (Verzi et al., 2015). Subterranean species were kept and tested at the animal facility of the University of Duisburg-Essen, Essen, Germany. Housing and testing of laboratory mice and rats took place at the Central Animal Laboratory, University Hospital Essen, Germany. Guinea pigs were housed and tested at the Department of Behavioural Biology of the University of Münster, Germany. Details on housing conditions were described for GMR in Begall et al. (2021) (with analogous husbandry for AMR, NMR and SC), for laboratory mice and rats in von der Beck et al. (2024), and for guinea pigs in Kaiser et al. (2022). Of each species, twelve adult individuals of both sexes were tested (six males, six females). Mean ± SD and ranges of body mass and age are given for each species in Table S1. While subjects from subterranean species were oftentimes markedly older than those from the epeigeic sample, we do not consider this an issue given their comparatively slow life histories and delayed senescence patterns (see e.g., Begall et al., 1999; Sahm et al., 2018). All tests were approved by the Landesamt für Natur, Umwelt und Verbraucherschutz NRW (approval number: 81-02.04.2023.A188).

Previous studies have stressed that the skin of the NMR is susceptible to evaporative water loss (Menon et al., 2019) and dry skin could critically bias test results. However, the skin of our subjects appeared silky and supple and did not display the flakey appearance typical of dry NMR skin (Delaney et al., 2021) so that we did not consider this an issue.

#### Procedure

Experiments were performed with a Cutometer® probe (Courage + Khazaka, Cologne, Germany). Prior to testing, a skin patch of about 2 cm x 2 cm was shaved on the ventral and dorsal side posterior to the rib cage in all furred subjects to enable the probe to make contact with the skin. In preparation for the tests, mice and rats were trained to keep still during handling (each individual was handled 2-3 times per week for approximately five minutes over a period of about four weeks). Such training was not necessary for the more docile guinea pigs. Subterranean rodents could not be effectively trained and had to be anaesthetized with isoflurane narcosis (UNO gas exhaust unit; flooding of the induction chamber with 5 % isoflurane; reduced to 2 % at the mask). Before starting the measurements, the cutometer base station (dual MPA 580) generated a negative pressure of 450 mbar, whereupon calibration of the probe took place. After successful calibration, the probe (MPA 580, aperture 2 mm) was gently pressed onto the (shaved) skin at a 90° angle. A single measurement required 6 seconds and was divided into a suction phase and subsequent relaxation phase: During the suction phase, which lasted for three seconds, the skin was sucked into the probe. In the following relaxation phase, the pressure dropped to 0 mbar to allow the skin to return to its original conformation. Optic sensors inside the probe measure the deformation of the skin (penetration depth into the probe) in 0.01 s intervals, so that a total of 600 values are generated during one measurement. For the dorsal and ventral skin, respectively, three valid measurements taken at adjacent skin areas (during which the probe was in airtight, uninterrupted contact with the skin, and thus no unintended drop in pressure during the suction phase occurred) were obtained for each individual. During data collection, ambient temperature was 22.5 ± 0.94 °C and relative air humidity was 43.3 ± 9.6 %. Test results are compiled in Table S5.

#### Statistics

In all calculations and visualizations, the offset value (the initial skin deformation caused by the pressure that the experimenter exerts by gently pressing the probe onto the skin) was taken into account by adding it to the respective parameter (Müller et al., 2018). Skin deformation curves (representing the penetration depth into the probe) and relevant parameters were generated automatically by the Cutometer® software (MPA CT plus, version 1.1.5.0). Parameters for the ventrum and dorsum were analyzed separately. We used LMMs to test for effects of ecology on the following parameters for dorsal and ventral skin, respectively: 1. maximum penetration depth into the probe (in mm) at the end of the suction phase, R0, indicating firmness of the skin, i. e. the lower R0, the firmer the skin; 2. regression distance (in mm) at the end of the relaxation phase, R8, reflecting the total recovery in the relaxation phase, i.e., the higher R8, the greater the skin’s ability to recover its initial conformation; 3. relaxation / suction ratio (in %), R2 (=R8/R0), the higher the value of this ratio the greater is the elasticity of the skin. Figure 2 visualizes the aforementioned parameters, our terminology for variables follows the one applied by the manufacturer of the Cutometer® system. The parameter R1 (R0-R8) which is generally considered to represent the ability of the skin to recover is not useful in interspecific comparisons if R0 values differ noticeably between the species.

**Figure 2:**
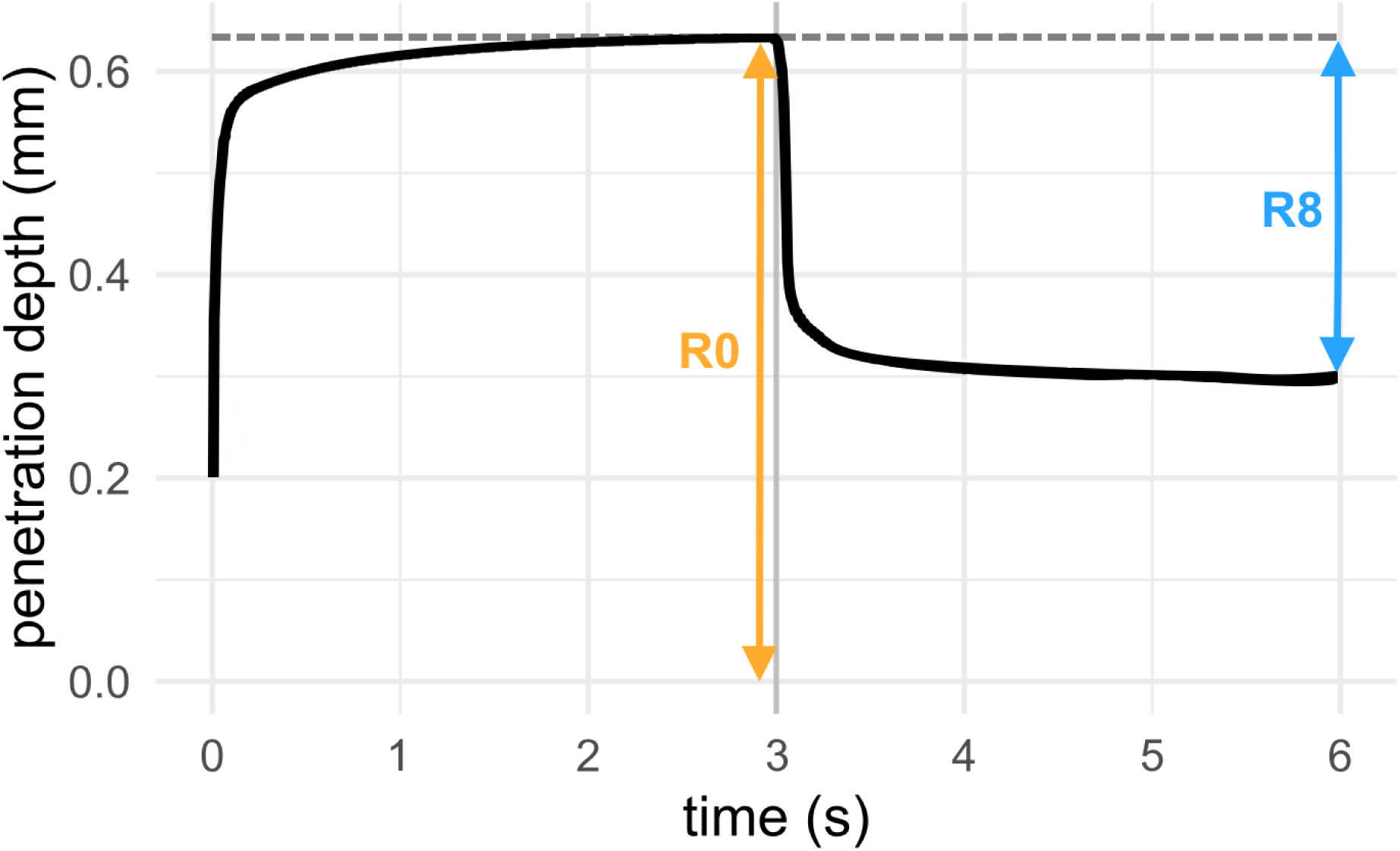
Idealized deformation curve (corrected by the offset value) from a single Cutometer® test. The offset value is 0.2 mm. R0 (orange arrow) indicates the maximum penetration depth into the probe at the end of the suction phase (at 3 seconds). R8 (blue arrow) indicates the recess of the skin at the end of the relaxation phase (at 6 seconds). R2 (= R8/R0) is derived from these measurements and represents the ratio of relaxation / suction.

Models for R0, R2, and R8 included ecology (epigeic/subterranean) and body mass (g) as fixed factors, as well as the interaction of said variables. Species as well as subject ID were additionally included as random factors (nested design: species/ID), resulting in the model formula R*x* ∼ ecology * body mass + (1|species / ID). Due to our small species sample (which prevented us from using phylogenetically-informed statistics; see e.g., Freckleton et al., 2002), the estimation of the models’ random effect variance may be imprecise. Nevertheless, we found our approach to be the most feasible to account for the non-independence of observations within species in our analyses. Model assumptions were checked using diagnostic plots and variance inflation factors were calculated to assess multicollinearity of fixed effects. Response variables were transformed either logarithmically or by square root to achieve normal distribution of model residuals.

### Uniaxial tensile testing of rodent skin

Although it has already been shown that HA has little effect on the tensile behavior of skin (Kronick & Sacks, 1994; Oxlund & Andreassen, 1980), we also aimed to test how excised skin flaps from the species we examined for elastic pliability respond to horizontal strain. Given methodological and sample-size related limitations, we chose to only include one incidental finding from these experiments here, namely the exceptional tensile resilience of NMRs skin when compared to the other species tested. Details on the testing procedure are included in Part III of the Supplementary File (Figs. S5-S6; Tables S3-S4). These results will be discussed alongside other findings on naked mole-rat skin biomechanics.

## Results

### Hyaluronan polymer size and distribution in tissues of Ansell’s mole-rat

For better contextualization, we report our findings on HA in AMR alongside the previously published results on NMR and guinea pig by del Marmol et al. (2021), who employed the same methodology as our study. AMR skin samples did not show any peculiarities regarding HA staining intensity or HA localization in any of the examined tissues. In general, these patterns appear to be conserved between AMR, NMR and guinea pig (Fig. 3). HA concentrations in skin and skeletal muscle were higher in the two bathyergids compared to the guinea pig (Table 1). In contrast, HA concentration in the kidney was higher in the guinea pig compared to AMR and NMR. Overall, HA concentrations in AMR tissues (Table 1) resembled that of the NMR, except for dermal hyaluronan, which was found to be higher concentrated in the skin of AMR samples.

**Figure 3:**
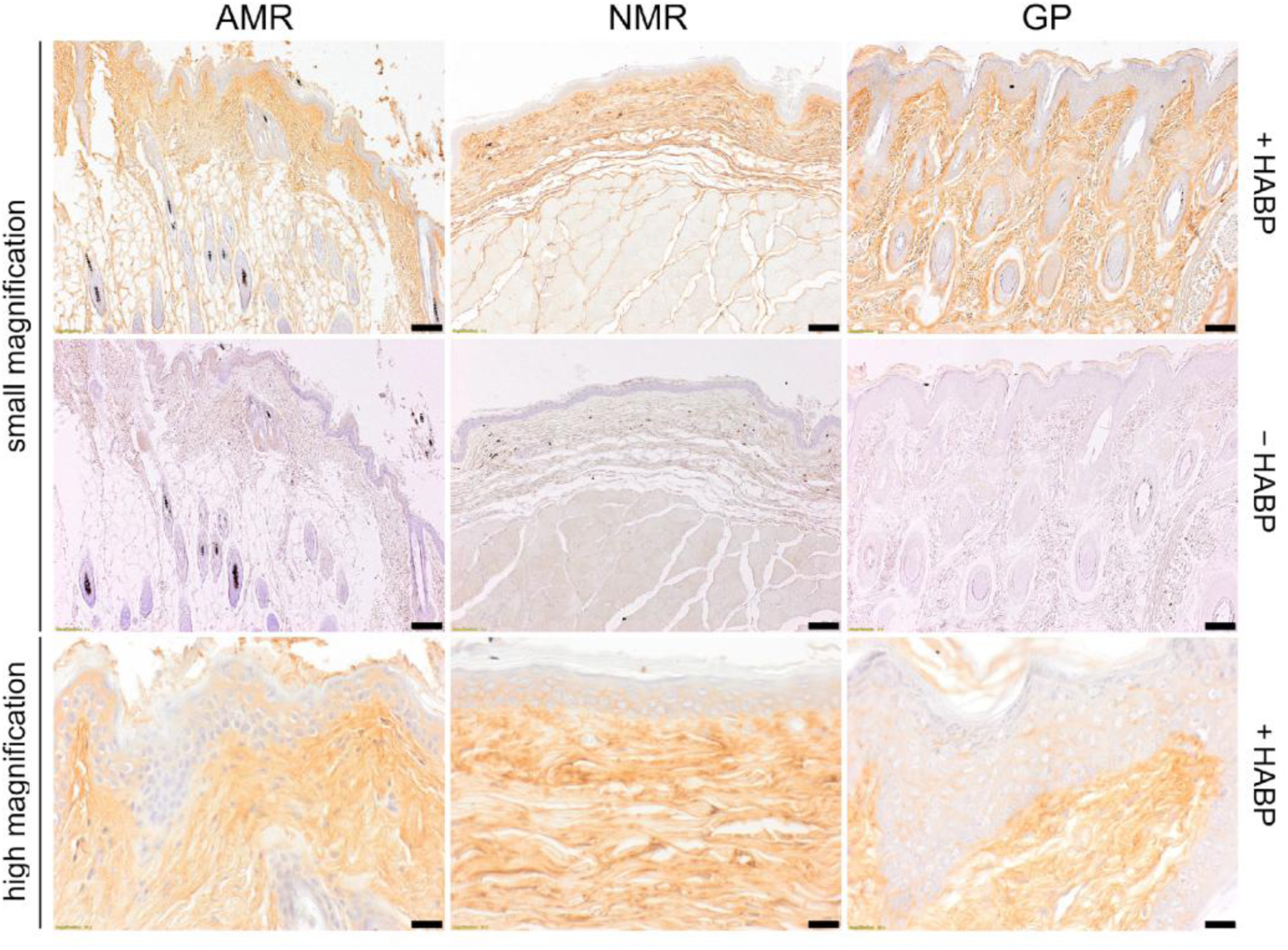
Histology of the skin in Ansell’s mole-rat (AMR) highlighting hyaluronan (HA), compared with previously published histologic images of tissues from the naked mole-rat (NMR), and guinea pig (GP). Tissues were stained following the same protocol and using HA binding proteins (HABP) followed by peroxidase detection. Positive signals are revealed in brown. Tissues are shown at small magnification with HABP (+HABP) and without (-HABP) as control (Scale bars, 100 µm). Tissues are shown at high magnification only when treated with HABP (scale bars, 20 µm). Images from NMR and GP tissue were retrieved from del Marmol et al. (2021).

**Table 1:**
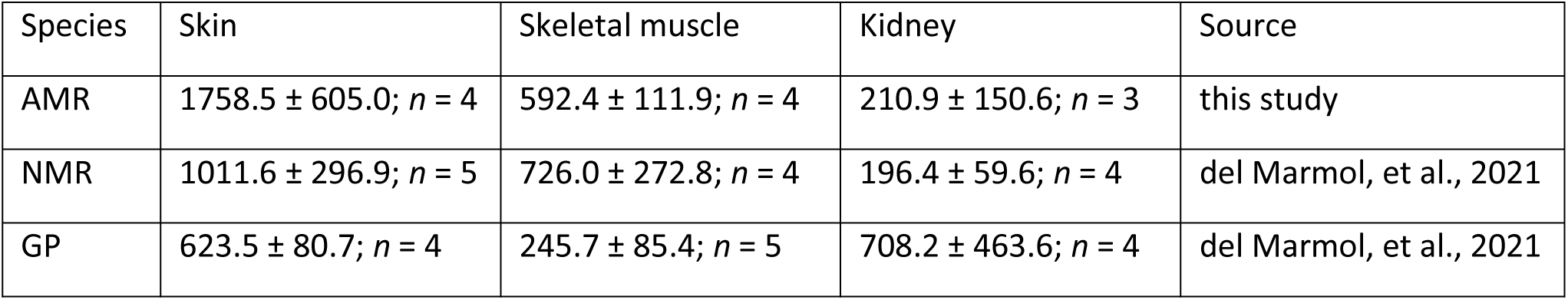
Hyaluronan concentration (ng/mg dry weight; mean ± SD) and numbers of individuals sampled in Ansell’s mole-rat (AMR), compared with naked mole-rats (NMR) and guinea pigs (GP) studied by del Marmol et al. (2021). Note that cross-study comparisons need to be undertaken with caution, given that separate assay runs of solid phase sandwich ELISA were performed.

There was no evidence for abundant HA of exceptional molecular mass (> 3 MDa) in the skin (Fig. 4) or any other tissue of the AMR (Fig. S2). Figure 4 illustrates the relative abundances of HA polymers of varying molecular mass in the sampled AMR skin compared to distributions in the NMR and the guinea pig (retrieved from del Marmol et al. 2021). In all sampled tissues, guinea pig HA tends to be shorter than that of the two bathyergid species, whose HA profiles are more similar to each other than either is to that of the guinea pig (see also Fig. S2). Still, the overlap between all three species is substantial.

**Figure 4:**
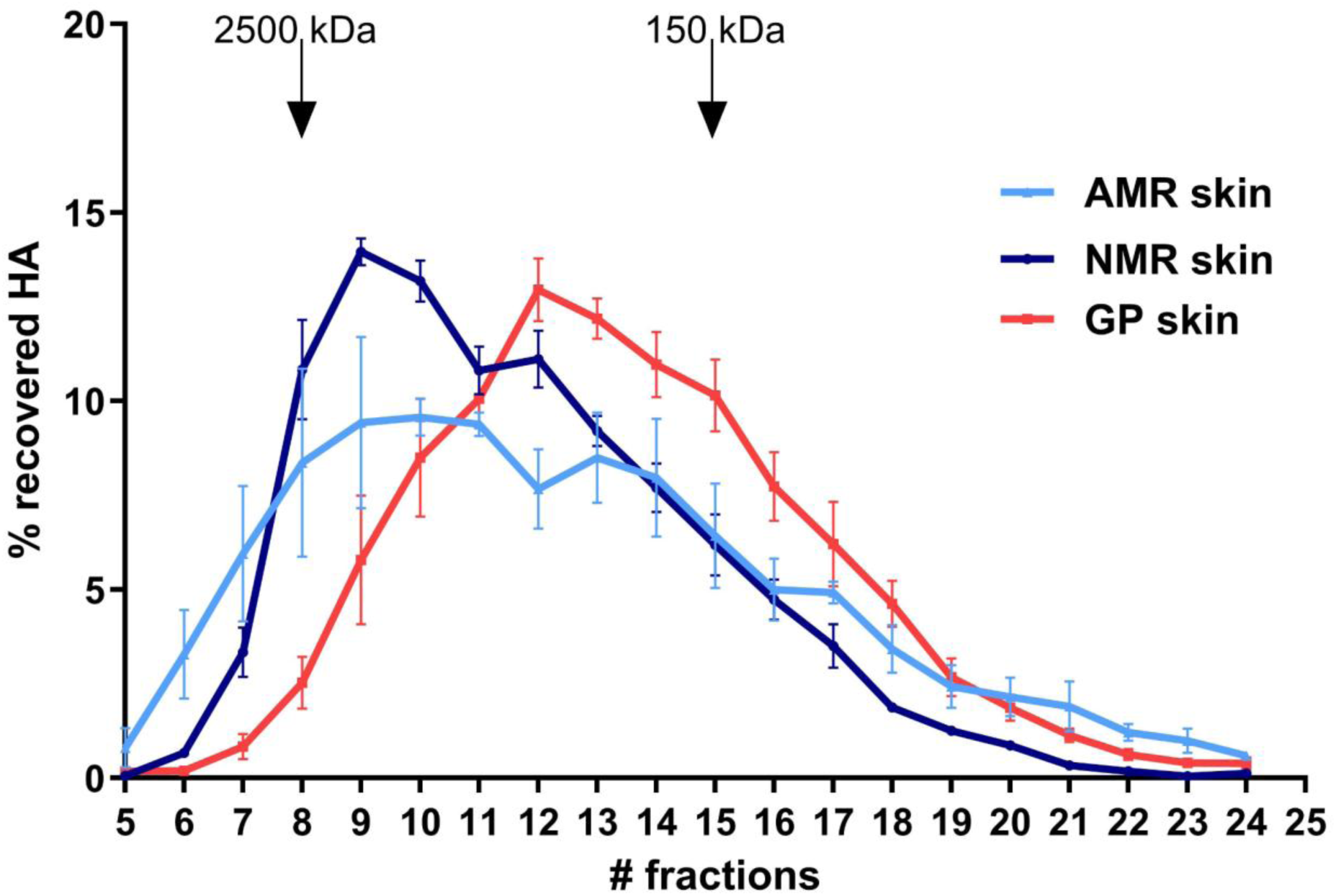
Hyaluronan (HA) molecular weight distributions from the skin of the Ansell’s mole-rat (AMR), determined by FPLC-SEC. Findings are compared to those of the naked mole-rat (NMR) and the guinea pig (GP). Data on NMR and GP were retrieved from del Marmol et al. (2021). Means and SEM are shown for each fraction. The elution peaks of two HA standards (2500 kDa and 150 kDa, respectively) are indicated by arrows.

HA polymers ranging between 150 kDa and 2500 kDa in size are the most abundant in AMR skin but the overall distribution of polymer sizes is broad and includes considerable amounts of both larger and smaller molecules (Fig. 4). In the NMR, dermal HA polymer sizes are similarly distributed (del Marmol et al., 2021) but polymers close to 2000 kDa are particularly abundant and those larger than 2500 kDa appear more poorly represented compared to AMR. Comparative HA size profiles for skeletal muscle and kidney are included in Fig. S2, the comparative histology of these tissues is shown in Fig. S3.

### Vertical deformation (Cutometer® tests)

Skin deformation curves (Fig. 5) revealed that the skin of the NMR is exceptionally stiff, which is reflected by low R0 values, indicative of high skin firmness. The NMR thus presents itself as an outlier in the dataset, considering both dorsal (Fig. 5a) and ventral measurements (Fig. 5b). We thus decided to run our statistical analyses on Cutometer® tests twice, including (Table 2) and excluding NMR data (Table S2). However, both sets of models were found to agree on which factors significantly influenced the studied parameters.

**Figure 5:**
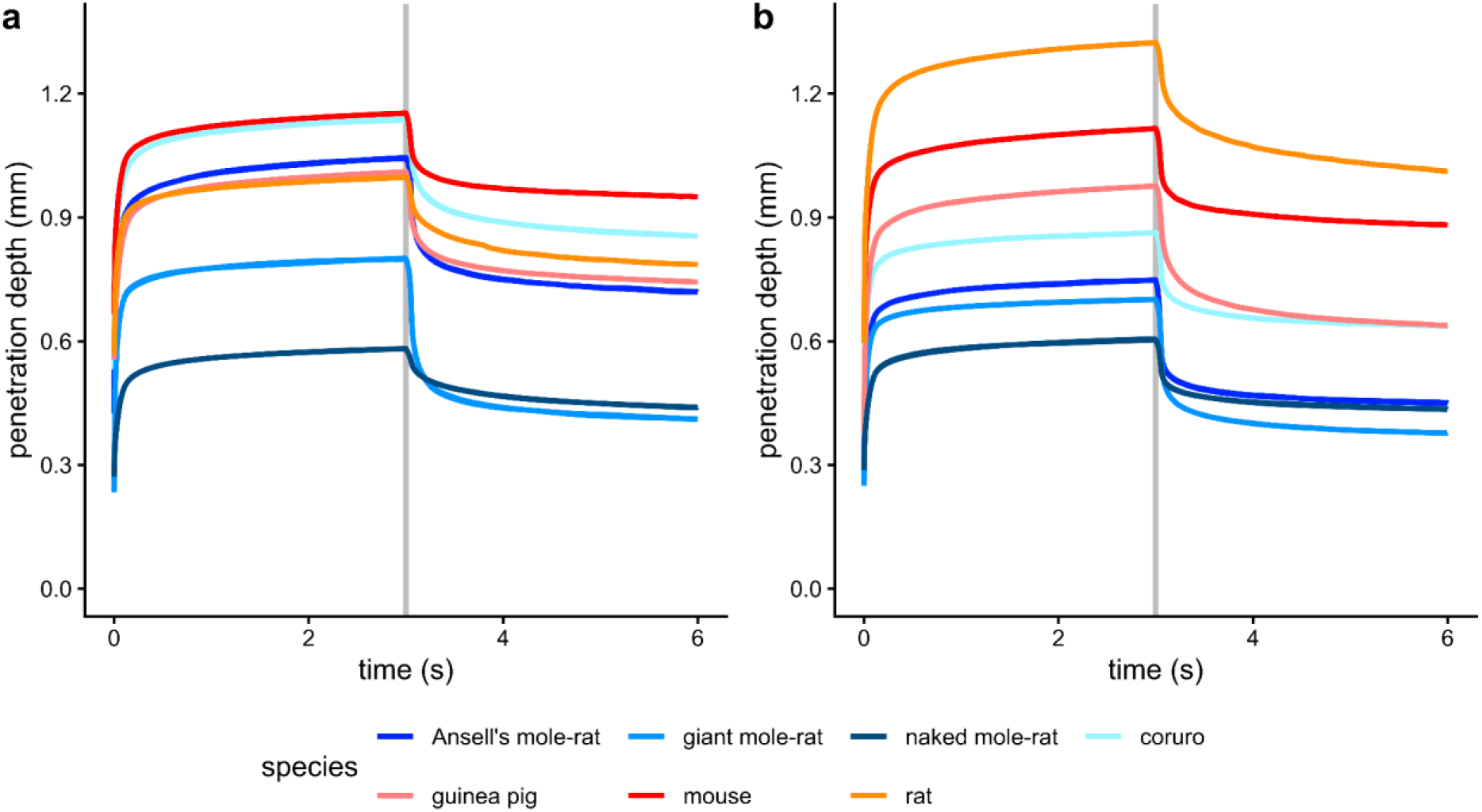
Species-level averaged skin deformation curves obtained by means of Cutometer® measurements at the dorsum (**a**) and ventrum (**b**). Measurements were corrected for the offset value with curves starting at the respective mean offset values. During measurements, suction was applied for three seconds, after which the skin was released to return to its initial conformation. Penetration depth indicates how far the skin was sucked into the probe. Grey lines indicate the end of the suction phase.

**Table 2:**
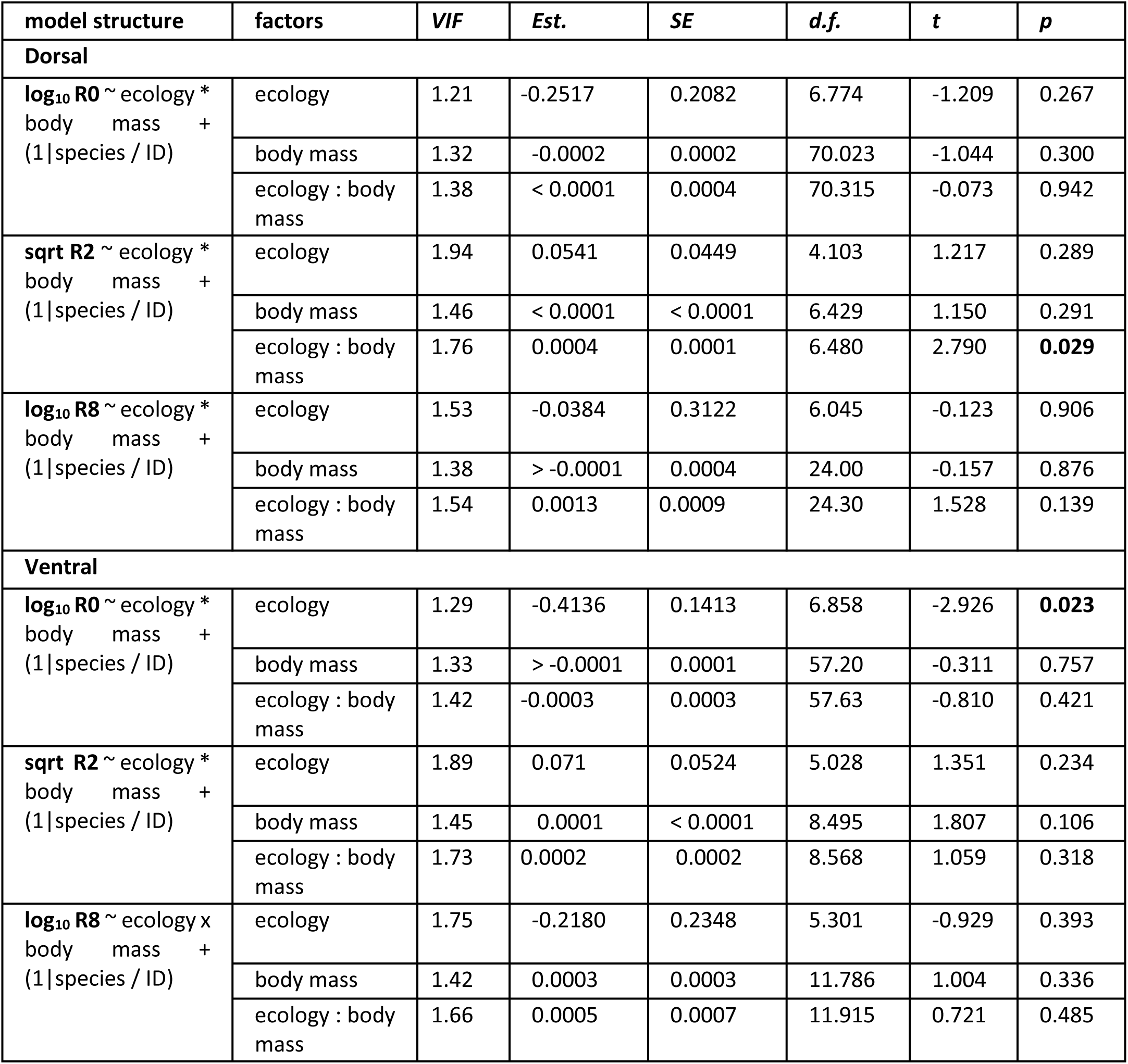
Results from linear mixed effect models on elasticity variables provided by Cutometer® measurements (dorsal and ventral regions were analyzed separately). Significant results are marked in bold. R0 indicates the maximum penetration depth at the end of the suction phase. R8 indicates the recess of the skin at the end of the relaxation phase. R2 is the ratio of relaxation / suction, with higher values indicating greater elasticity. For the factor ecology, “subterranean” is always the reference. Est.: estimate, SE: standard error, VIF: variance inflation factor. Respective transformation (log_10_, square root) of the response variable is indicated and estimates were not back-transformed.

Subterranean rodent skin was not found to be overall more elastic than that of epigeic forms (Table 2). In fact, only very few significant differences in skin pliability could be found between the two ecological groups and species differences within said groups were at times pronounced (Fig. 6). Skin deformation curves for the ventral region (Fig. 5b, Fig. S4) indicate greater skin firmness in all subterranean species when compared to epigeic ones, as penetration depth was found to be low overall. In accordance, the respective LMM indicates that R0 values for the ventral side are significantly lower for subterranean species compared to epigeic ones, regardless of whether the NMR is included (Fig. 6, Tab. 2; *p =* 0.023) or not (Tab. S4; *p* = 0.030). For the dorsal region, a significant effect of ecology did not emerge in either model (Tab. 2; Tab. S4). We also recovered a significant interaction between ecology and body mass for R2 values (Fig. 6, Tab. 2, Fig. S4) at the dorsum (*p* = 0.029) but not the ventrum (*p* = 0.318), both with and without the NMR present in the species sample (Tab. S4). Thus, the effect of body size differs between the epigeic and subterranean groups in that regard. While body size is positively correlated with dorsal skin R2, a proxy for skin elasticity, in the underground-dwelling taxa, this is not the case in epigeic ones.

**Figure 6:**
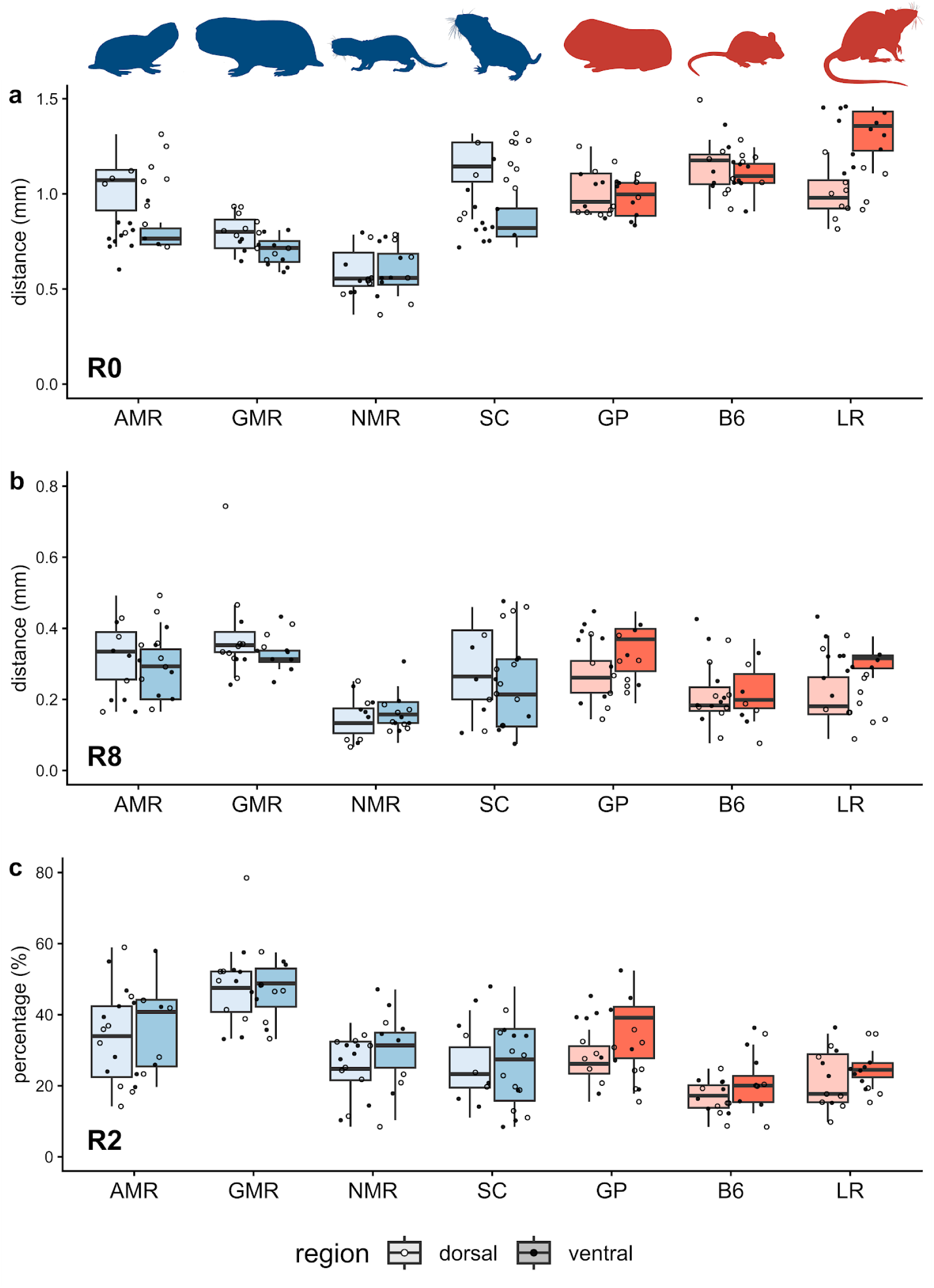
Boxplots illustrating biomechanical skin properties determined via Cutometer® tests. Subterranean species are shown in blue shades, epigeic ones in red shades. Points represent mean values of each tested individual. Closed points correspond to dorsal measurements, open ones to ventral ones. **a**: R0 values (maximum amplitude of skin deformation at the end of the suction phase). **b**: R8 values (maximum regression distance of skin at the end of the relaxation phase). **c**: R2 values (R8/R0; relaxation distance / suction distance). AMR: Ansell’s mole-rat, B6: C57BL/6 laboratory mouse, GMR: Giant mole-rat, GP: Guinea pig, LR: LEW/Crl laboratory rat, NMR: Naked mole-rat. Silhouettes derive from PhyloPic (https://www.phylopic.org/).

### Uniaxial tensile testing: Unusual tensile properties of naked (mole-rat) skin

When exploring tensile properties of rodent skin (Suppl. Information Part III), we noticed that the skin of the naked mole-rat is unusual in enduring extreme stresses before perforating (Fig. 7, Table S3; mean max. stress - dorsal, *n* = 5: 9.3 MPa; ventral, *n* = 3: 8.6 MPa). We hypothesized that these effects could be related to the lack of pelage and related structural differences in the naked mole-rat dermis and tested this idea by examining skin samples of a naked lab mouse strain (NMRI-nu). The dorsal skin samples of naked lab mice (*n* = 2) could endure average maximum stresses (5.1 ± 1.41 MPa) more than twice as high than those of C57BL/6 mice (2.23 ± 0.06 MPa), with the yield stress also being elevated in the naked strain (0.55 ± 0.13 MPa) compared to the C57BL/6 mice (0.32 ± 0.12 MPa). The scarcity of available samples precluded statistical comparisons as well as a sufficient characterization of ventral skin regions (Suppl. Information Part III).

**Figure 7:**
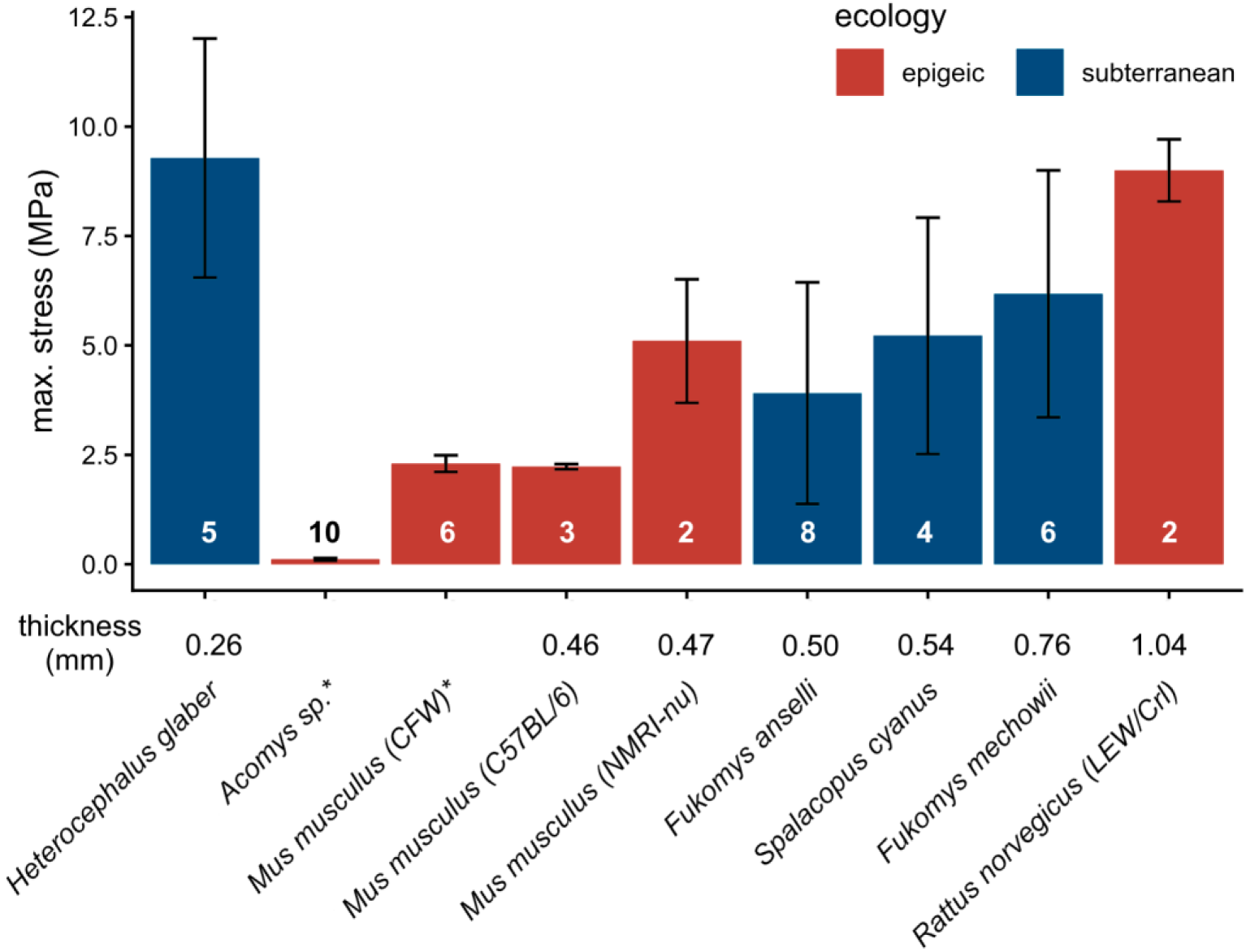
Bar plots showing mechanical stress tolerated by the dorsal skin of various rodents, including three lab mouse strains. Groups are arranged from left to right in order of increasing skin thickness. Sample sizes are indicated by numbers plotted on respective bars. Numbers below bars indicate mean skin thickness in mm (measured with calipers on excised skin flaps), error bars indicate standard deviations. Statistical comparison was not feasible due to limited sample sizes. Note that NMR and nude mice (NMRI-nu) exhibit notably resilient skin. All samples, except for those from C57BL/6 mice were frozen and thawed before testing. *Data derived from Seifert et al. (2012), no data on mean skin thickness were provided. Samples sizes and methodological details are included in Part III of the Suppl. Information.

## Discussion

### Hyaluronan in African mole-rats

The HA polymer size distribution in the tissues of the AMR was found to be similar to that of the NMR (Fig. 4) (del Marmol et al., 2021), with the skin of the former showing a greater abundance of very long (2500 kDa) HA molecules. Both bathyergids displayed longer dermal HA polymers than the guinea pig and also laboratory mice (del Marmol et al., 2021). Whereas absolute amounts of dermal HA in the bathyergids also appear to be notably higher than those of epigeic species studied with ELISA-like assays (Bourguignon & Flamion, 2016; del Marmol et al., 2021), cross-study comparisons of absolute HA levels should be treated with caution. Interestingly, differences in hyaluronan binding proteins (HABP) fluorescence intensity also indicate higher levels of HA polymers in the bathyergid dermis compared to that of the guinea pig (Zhao et al., 2023), supporting the assumption that there are genuine differences between the species in this respect. Applying the same method, Zhao et al. (2023) also reported higher concentrations of HA in the kidney of bathyergids (NMR and Damaraland mole-rat), especially the NMR, when compared to the guinea pig. The data of del Marmol et al. (2021) do not corroborate this finding since HA concentration was lower in the kidney of the NMR compared to the guinea pig and our findings suggest a similar pattern for AMR. In any case, AMR tissues were not found to contain exceptionally long HA polymers (> 3 MDa) in notable amounts, questioning its ubiquitous presence in subterranean mammals (Braude et al., 2021; del Marmol et al., 2021). This finding is at odds with the data reported on other bathyergids in the studies by Tian et al. (2013) and Zhao et al. (2023), which have previously been criticized for the methods used to identify said molecules (del Marmol et al., 2021; see Introduction). Systematic comparisons of the different methodological approaches employed are required to better delineate limitations of both and how those might affect the data yielded from them.

However, all available studies, including our own, agree that bathyergids display higher HA concentrations and at least moderately elongated HA polymers in their skin than epigeic laboratory rodents and it is tempting to speculate about potential functions of these traits. Our results as well as previous findings suggest that such differences in HA phenotypes do not impact skin elasticity, when horizontally (Kronick & Sacks, 1994; Oxlund & Andreassen, 1980) or vertically applied strain (this study) is concerned. However, this might be different for skin compression. Because longer, highly concentrated HA polymers can bind water more effectively, they create a more viscous and resilient extracellular matrix, which in turn might be able to shield capillaries and other microstructures in the skin against damage from friction experienced during burrowing. In the NMR in particular, other skin modifications are known which might ameliorate friction-related stresses on skin tissue. These include a thickened epidermis as well as extensive macroscopic folding of the skin, buffering mechanical stresses to compensate for the lack of pelage (Sokolov, 1982; Tucker, 1981; Daly & Buffenstein, 1998). We want to note here that the pronounced stiffness of the skin we recovered for the NMR by aid of the Cutometer® probe, might to some extent be a side effect of its epidermal specializations. Kulaberoglu et al. (2019) studied the material properties of purified NMR HA and found it folds into complex and elastic 3D conformations, which might contribute to making it an efficient mechanical buffer *in vivo*. To our best knowledge, an effect of HA polymer length on skin compression properties has not been experimentally demonstrated yet, but it is evident that HA contributes to buffering such stresses (Tregear, 1969). Comparative studies on the behavior of skin under compression in African mole-rats and other small mammals would be essential to test this idea.

Regardless of whether dermal HA plays a role in subterranean adaptation, the upregulation and elongation of HA polymers across various tissues in bathyergids, but also other subterranean mammals (Zhao et al., 2023; Fig. 4 and Fig. S2) calls for additional explanations beyond those concerning skin characteristics. For instance, both NMRs and species of the genus *Fukomys*, to which the AMR belongs, display notable longevity and cancer resistance (in both genera, several cases of cancer have nevertheless been described - Delaney et al., 2021; Sakai et al., 2025; Sura et al., 2010). For the NMR, HA polymer length and abundance have been hypothesized to contribute to these traits, by granting early contact inhibition to limit cell proliferation (Tian et al., 2013). Nevertheless, whether HA might also be involved in *Fukomys* cancer resistance remains to be shown (Suzuki et al., 2025) and so far there is no indication that it is crucial for tumor suppression in any other subterranean rodent (such as the cancer-resistant blind mole-rat - Manov et al., 2013). Thus, a consistent link between elongated HA polymers and cancer resistance remains highly speculative.

High concentrations of elongated HA polymers have also been suggested to be involved in tolerating hypoxic conditions (Zhao et al., 2023), which are commonly encountered by subterranean mammals. There is evidence that hypoxia is associated with increased damage by reactive oxygen species generated in the mitochondria (Solaini et al., 2010) and in line with that, subterranean mammals display derived mechanisms to effectively reduce oxidative stress (Munro et al., 2019; Schülke et al., 2012). Interestingly, there is evidence that HA secreted by NMR fibroblasts reduces intracellular oxidative damage via the HA-binding CD44 receptor (Takasugi et al., 2020). These findings require replication and whether they can be translated to other subterranean species remains unclear. NMRs express high-levels of CD44, a trait not shared among subterranean mammal lineages (Takasugi et al., 2023). Still, tolerance to hypoxia should be considered as a potential driver for derived HA phenotypes in underground-dwelling mammals.

### Skin elasticity

We failed to recover evidence for consistently increased skin elasticity in subterranean compared to epigeic forms using *in vivo*-cutometry, challenging the idea that these groups should clearly deviate from one another in this trait (Tian et al., 2013; Zhao et al., 2023). This is in line with research identifying the extracellular fiber network of the skin as the primary determinant of skin elasticity in mammals (Silver et al., 2003), not mucopolysaccharides such as HA. Overall, we recovered only a few statistically significant differences between burrowing and epigeic rodents, but their interpretation is not straight forward. For instance, it would be intuitive to hypothesize that the firm ventral skin found across subterranean species is an adaptive trait countering friction when moving through underground tunnels. In that case, however, firmer dorsal skin would appear beneficial as well (Kimani, 2013) but is not present in subterranean species except for the NMR (Fig. 5). It can be argued that firmer ventral skin in furred subterranean species is a non-adaptive side effect of a lower hair density on the ventrum. In both African mole-rats and the coruro, experimental data suggest that the venter acts as a thermal window for heat exchange (Vejmělka et al., 2021), which is facilitated by thinner pelage. For instance, hair density in the giant mole-rat is approximately four times greater on the back (ca. 11154 x cm^−2^) compared to the belly (ca. 2753 x cm^−2^ – Šumbera et al., 2007), while in the similarly-sized Norway rat, hair density is lower on the dorsum (ca. 5530 x cm^−2^) than on the ventrum (7705 x cm^−2^ – Sokolov, 1982; data for winter pelage). In thin-skinned small mammals, the relative portion of the adnexa, especially hair follicles, compared to total skin volume is high. However, these structures occupy space which would otherwise be filled by the dermal collagen fiber network that underlies the skin’s tensile strength. Thus, skin firmness should be inversely correlated with hair density in these animals. We will revisit this principle when discussing the tensile strength of naked mole-rat skin later on.

Given that both the tested epigeic and subterranean species showed pronounced variation within their ecological groups, our species dataset needs to be expanded before firm conclusions about potential links between ecology and skin elasticity can be drawn. Given the limited availability of species to investigate, the subterranean rodents tested here all were rather closely related tooth-digging hystricomorphs and the epigeic ones derive from only two very distant and poorly represented branches of the rodent order, murids and caviids.

Unfortunately, causes for interspecific differences in skin biomechanics are not well understood. This makes it difficult to interpret, for instance, the notably high skin elasticity we found in giant mole-rats. When the skin experiences minor deformation and strain below the yield point, as was induced by the Cutometer® probe, dermal elastin fibers act as the main load bearing component and also enable its rebound into the initial conformation (Silver et al., 2003). A denser network of elastin fibers would thus be expected to result in increased elasticity (Debelle & Tamburro, 1999). In domestic rats and guinea pigs, elastin abundance is greater in ventral compared to dorsal skin (Meyer et al., 1994). In line with this, we found elasticity of ventral skin (as approximated by R2) to be greater than that of dorsal skin in these species. In fact, elasticity was significantly more pronounced on the ventrum at the level of our total sample. Previous studies on subterranean rodents have noted dense elastin fiber networks in the skin of AMR (Hesselmann, 2010) and NMR (Kimani, 2013; Sokolov, 1982) as well as in burrowing spalacid and cricetid rodents (Sokolov, 1982). Interestingly, this pattern does not appear universal in subterranean mammals, since the elastic fiber system is only weakly developed in talpid moles and pocket gophers (Giacometti & Machida, 1965; Starcher, Aycock, & Hill, 2005). On the other hand, dense elastic fiber systems can also occur in epigeic species. For instance, laboratory rats display dermal elastic fiber networks that resemble those of bathyergid mole-rats (Hesselmann, 2010), yet their skin is notably less elastic. Again, a dichotomy between subterranean and epigeic forms is not apparent. The structure and potential allometric constraints on the elastic system of small mammals require further study.

We did not investigate horizontal skin elasticity nor skin looseness (Fig. 1). As previously discussed, available experimental data do not suggest that HA polymers impact horizontal skin elasticity to notable extents (Kronick & Sacks, 1994; Oxlund & Andreassen, 1980). Hence, we do not expect that subterranean mammal skin can be stretched further than that of epigeic mammals of comparable skin thickness, before accruing lasting damage. However, to conclusively test this hypothesis, freshly excised or appropriately stored skin flaps of relevant taxa would need to be subjected to controlled uni- or biaxial tensile tests to determine elastic properties such as yield stress and strain of the tissue (analogous to tests focusing on maximum stress reported herein). Previous work on mammal skin’s elastic behavior has shown great variation between samples of the same species (e.g., Caro-Bretelle et al., 2015), making it challenging to acquire adequate sample sizes for meaningful comparative analyses. We are not aware of any such experiments with a focus on burrowing mammal species.

Whether subterranean mammals exhibit increased skin looseness, a phenomenon that may have been confused with skin elasticity in the past, remains to be shown but will be difficult to quantify. In any case, we want to emphasize that it does not appear feasible that HA may contribute to an increase in looseness. This trait is primarily dictated by the development of the subcutis, which attaches the dermis to the underlying musculature via collagenous sheets that are anchored by elastin fibers (Kawamata et al., 2003; Menton et al., 1978). To enable skin looseness, these subcutaneous collagen sheets need to be lubricated so that they can gently slide past each other. HA contributes to this lubrication, but there is evidence that an increase in HA concentration and molecular mass actually impairs rather than aids mobility in such connective tissue fibrillary networks (Cowman et al., 2015).

Finally, we want to briefly discuss the remarkable mechanical resilience of NMR skin, which can endure maximum stresses that vastly exceed those that can be applied to the skin of small mammals of comparable body size and skin thickness. Our preliminary finding that the skin of similar-sized naked lab mice also shows elevated tensile strength suggests that the absence of fur in both groups contributes to this trait (Fig. 7). As discussed above, hair follicles occupy space in the dermis that can otherwise be filled by collagen-rich extracellular matrix, which in turn should increase the skin’s tensile strength. The impact that hair follicles can have on the mechanical resilience of small mammal skin is impressively exemplified by some species of African spiny mice (genus *Acomys*). These rodents have extremely fragile skin and exhibit substantially enlarged hair follicles, which most likely cause the low tensile strength of their dermis (Seifert et al., 2012; Fig. 7). Hence, the absence of pelage in the NMR and naked lab mice may have the opposite effect: The lack of hair follicles should allow for a more extensive dermal collagen fiber network and thus increased tensile strength. Our observations are in line with this reasoning. Given that, we suggest that the remarkable tensile strength of the NMR is an evolutionary byproduct of its nakedness and does not entail an adaptive value.

### Conclusions

We experimentally tested the hypothesis that greatly elongated HA polymers evolved in subterranean mammals to increase skin elasticity. Our results do not suggest that such molecules occur to notable extents in Ansell’s mole-rat. We could also have not found consistent differences in skin elasticity between subterranean and epigeic rodents based on *in vivo* cutometry. Nevertheless, these findings should be viewed as preliminary given the limitations of our species sample. The skin of the NMR stood out from our sample due to its great firmness and was also shown to exhibit high tensile strength, which could relate to the loss of pelage in this species. We argue that increases in HA polymer size do not enhance skin elasticity or looseness, which might have been confused with skin elasticity in the past. However, we show that not only NMRs but also AMRs express moderately elongated HA polymers across different tissues, a trait that might have evolved in the context of oxidative stress protection. It also appears feasible that longer and more abundant HA polymers might protect the skin from friction-induced compression damage. These hypotheses should be experimentally addressed by future studies.

## Supporting information

Supplementary File (Figs. S1 - S6; Tabs. S1 - S4)

## List of abbreviations

AMR: Ansell’s mole-rat
B6: laboratory mouse (C57BL/6)
DA: Dalton
FPLC-SEC: Fast Protein Liquid Chromatography-Size Exclusion Chromatography
GMR: giant mole-rat
GP: guinea pig
HA: hyaluronan
HABP: hyaluronan binding protein
LR: laboratory rat (Lewis strain)
NMR: naked mole-rat
SC: *Spalacopus cyanus*

## Acknowledgements

We are indebted to Courage + Khazaka (Cologne, Germany) for lending us the Cutometer® testing device, especially the help by Gabriele Uhl is highly appreciated. We thank Gero Hilken (Central Animal Laboratory, University Hospital, Essen, Germany) and Susanne Holtze (Department of Reproduction Management, Leibniz Institute for Zoo and Wildlife Research, Berlin, Germany) for the provision of crucial skin samples used for horizontal straining. We are grateful to Sylvia Kaiser (Department of Behavioural biology, University of Münster) for providing guinea pigs, and Gero Hilken for providing laboratory rats and mice for experiments. This study was partially funded by the German Society for Mammalian Biology (DGS e.V.). We thank an anonymous reviewer for constructive comments on the text which significantly improved the manuscript.

