## Supplementary File (Figs. S1 - S6; Tabs. S1 - S4) for "Hyaluronan and skin elasticity: Are subterranean mammals special?"

**Supplementary Information**

**Part I: Supplementary figures and data on hyaluronan polymers**

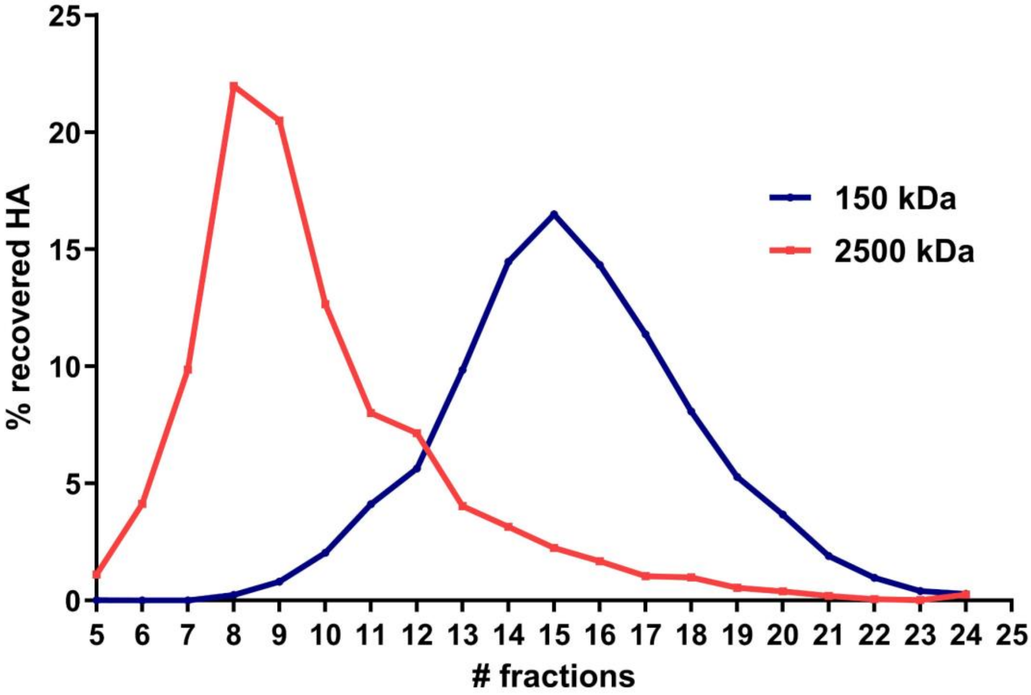


**Supplementary Figure S1**: Elution profiles of hyaluronan (HA) polymers for commercial HA markers (SelectHA; Hyalose LLC; SanDiego, CA, USA) used in this study.

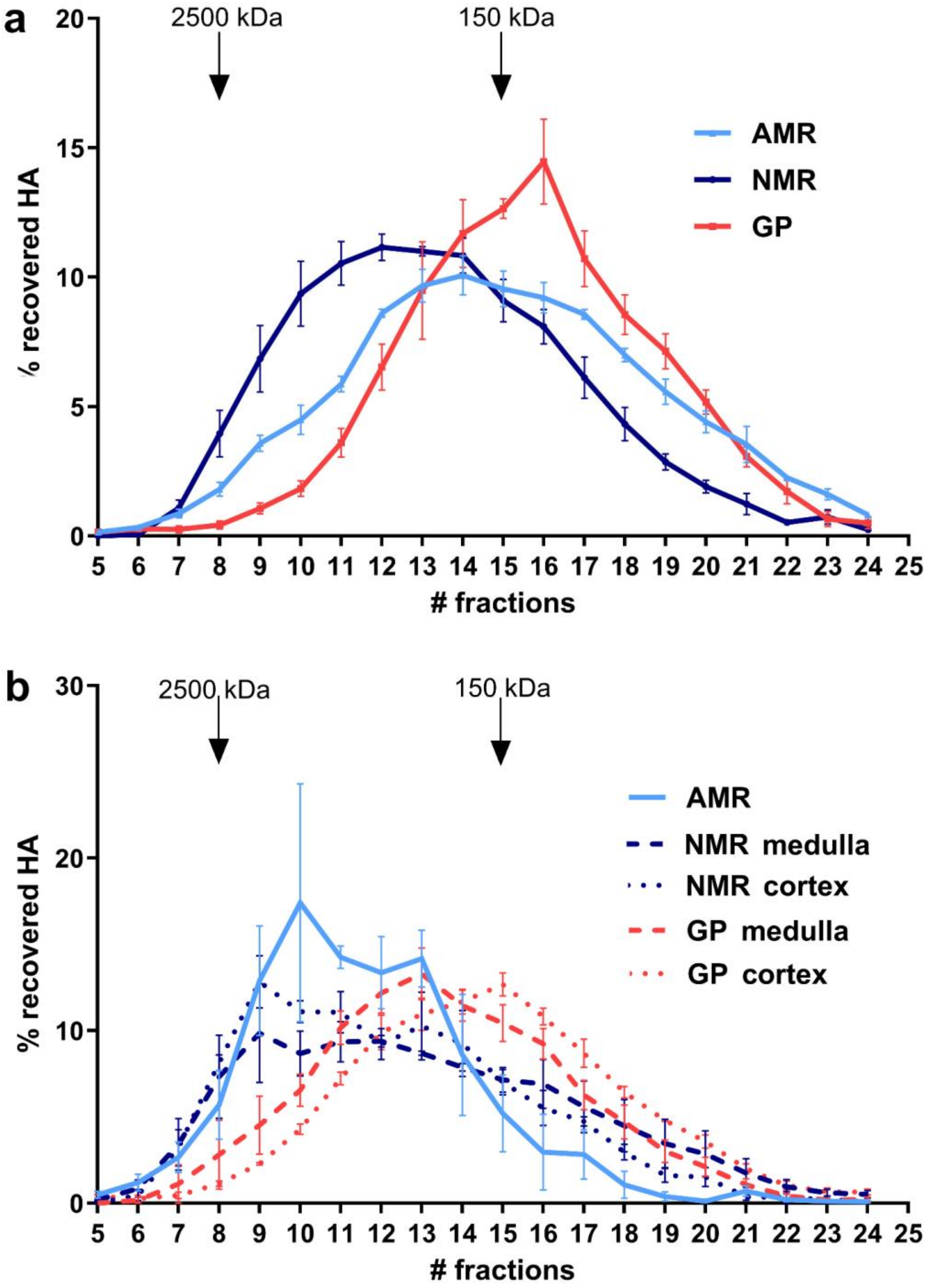


**Supplementary Figure S2**: Hyaluronan (HA) molecular weight distributions in the skeletal muscle (**a**) and kidney (**b**) of the Ansell's mole-rat (AMR) compared to those of the naked mole-rat (NMR) and the guinea pig (GP). Note that we could not partition kidney medulla and cortex for measurements of HA polymer size distribution in AMR. For the NMR and GP kidney results for medulla and cortex are separately presented. Both subterranean species tend to display a greater abundance of long polymers than the guinea pig. In skeletal muscle, AMR HA polymers sizes are shifted to lower molecular masses compared to the NMR. The relative abundance of renal HA polymers ranging between 150 kDa and 2500 kDa is notably higher in the AMR compared to the NMR, but lower for polymers smaller than 150 kDa. The AMR kidney profile displays two peaks, which might represent differences in HA polymer size distributions between the renal cortex and medulla, respectively.

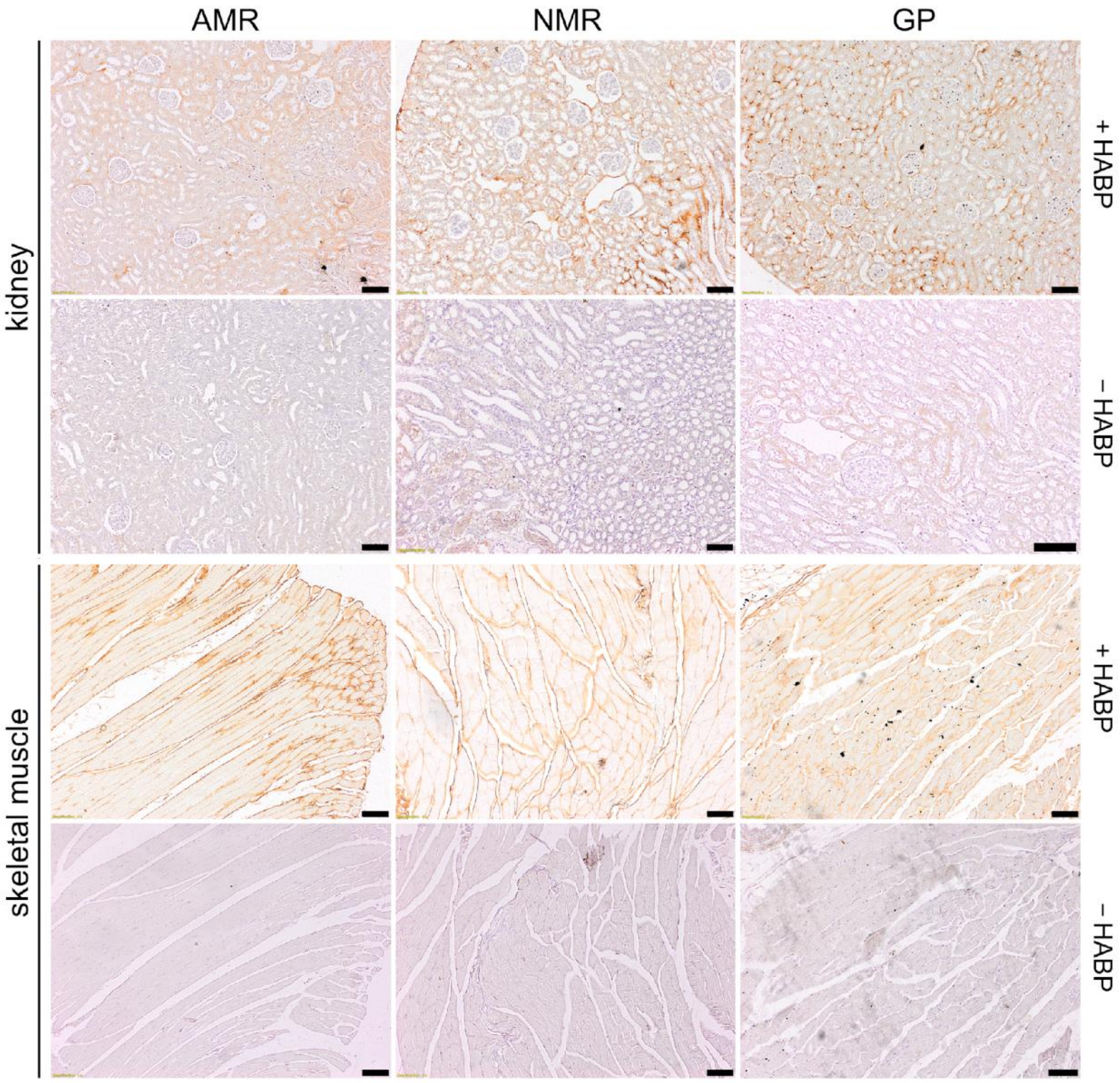


**Supplementary Figure S3**: Histology of the kidney and skeletal muscle in Ansell's mole-rat (AMR), naked mole-rat (NMR), and guinea pig (GP). Tissues were stained using HA binding protein followed by peroxidase detection. Positive signals are revealed in brown. Tissues are shown at small magnification with HABP (+HABP) and without (-HABP) as control (scale bars, 100 µm). Images from NMR and GP tissues were retrieved from del Marmol et al. (2021).

**Part II: Supplementary tables and figures on elastic pliability**

**Supplementary Table S1**: Mean ± SD and range (in parentheses) of body mass (g) and age (days) for seven tested rodent species used for measurements of skin elasticity (*n* = 12 for each species)

| **Species** | **Body mass (g)** | **Age (days)** |
|---|---|---|
| *Cavia porcellus* | 728 ± 208 (424.9-1054) | 134 ± 66 (52-235) |
| *Fukomys anselli* | 74.7 ± 21.4 (51.1-113.1) | 732 ± 242 (312-1408) |
| *Fukomys mechowii* | 371 ± 116 (216.2-622.7) | 1830 ± 1088 (707-3468) |
| *Heterocephalus glaber* | 38.8 ± 9.4 (24.6-59.3) | 1168 ± 640 (286-1831) |
| *Mus musculus* | 26.3 ± 5.2 (20.5-33.8) | 102 ± 22 (81-131) |
| *Rattus norvegicus* | 298 ± 91 (204-428) | 131 ± 32 (83-173) |
| *Spalacopus cyanus* | 119 ± 16.7 (101.4-149.5) | 885 ± 17 (201-2822) |

**Supplementary Table S2**: Results from linear models on elasticity variables provided by Cutometer® tests without data for naked mole-rats (dorsal and ventral regions were analyzed separately). Significant results are marked in bold. R0 indicates the maximum penetration depth at the end of the suction phase. R8 indicates the recess of the skin at the end of the relaxation phase. R2 is the ratio of relaxation / suction, with higher values indicating greater elasticity. For the factor ecology, “subterranean” is always the reference. Est.: estimate, SE: standard error, VIF: variance inflation factor, Est: estimate, SE: standard error. Respective transformation ($\log_{10}$, square root) of the response variable is indicated and estimates were not back-transformed.

| **Model** | **Factors** | **VIF** | **Est.** | **SE** | ***d.f.*** | ***t*** | ***p*** |
|---|---|---|---|---|---|---|---|
| **Dorsal** | | | | | | | |
| **$\log_{10}$ R0** ~ ecology * body mass + (1\|species/ID) | ecology | 1.48 | -0.0545 | 0.097 | 5.17 | -0.561 | 0.598 |
| | body mass | 1.30 | -0.0002 | 0.0001 | 30.53 | -1.796 | 0.082 |
| | ecology : body mass | 1.57 | -0.0003 | 0.0002 | 40.91 | -1.122 | 0.268 |
| **sqrt R2** ~ ecology * body mass + (1\|species/ID) | ecology | 1.48 | 0.0574 | 0.0577 | 4.42 | 0.995 | 0.371 |
| | body mass | 1.30 | < 0.0001 | <0.0001 | 17.67 | 0.291 | 0.775 |
| | ecology : body mass | 1.57 | 0.0004 | 0.0002 | 23.55 | 2.385 | **0.025** |
| **$\log_{10}$ R8** ~ ecology * body mass + (1\|species/ID) | ecology | 1.96 | 0.2872 | 0.1717 | 2.13 | 1.673 | 0.229 |
| | body mass | 1.33 | 0.0001 | 0.0002 | 3.76 | 0.600 | 0.583 |
| | ecology : body mass | 1.89 | 0.0007 | 0.0005 | 4.65 | 1.295 | 0.256 |

| | | | | | | | |
|---|---|---|---|---|---|---|---|
| **Ventral** | | | | | | | |
| **log10 R0** ~ ecology * body mass + (1\|species/ID) | ecology | 1.31 | - 0.3084 | 0.1069 | 5.65 | -2.884 | **0.030** |
| | body mass | 1.29 | > -0.0001 | 0.0001 | 73.13 | -0.433 | 0.666 |
| | ecology : body mass | 1.46 | -0.0004 | 0.0002 | 93.92 | -1.795 | 0.076 |
| **sqrt R2** ~ ecology * body mass + (1\|species/ID) | ecology | 1.54 | 0.0745 | 0.0611 | 5.27 | 1.220 | 0.274 |
| | body mass | 1.30 | < 0.0001 | <0.0001 | 24.54 | 1.383 | 0.179 |
| | ecology : body mass | 1.61 | 0.0001 | 0.0002 | 32.73 | 0.900 | 0.375 |
| **log10 R8** ~ ecology * body mass + (1\|species/ID) | ecology | 1.62 | -0.0604 | 0.2161 | 5.25 | -0.279 | 0.791 |
| | body mass | 1.31 | 0.0003 | 0.0003 | 19.09 | 1.046 | 0.309 |
| | ecology : body mass | 1.66 | 0.0001 | 0.0006 | 25.20 | 0.209 | 0.836 |

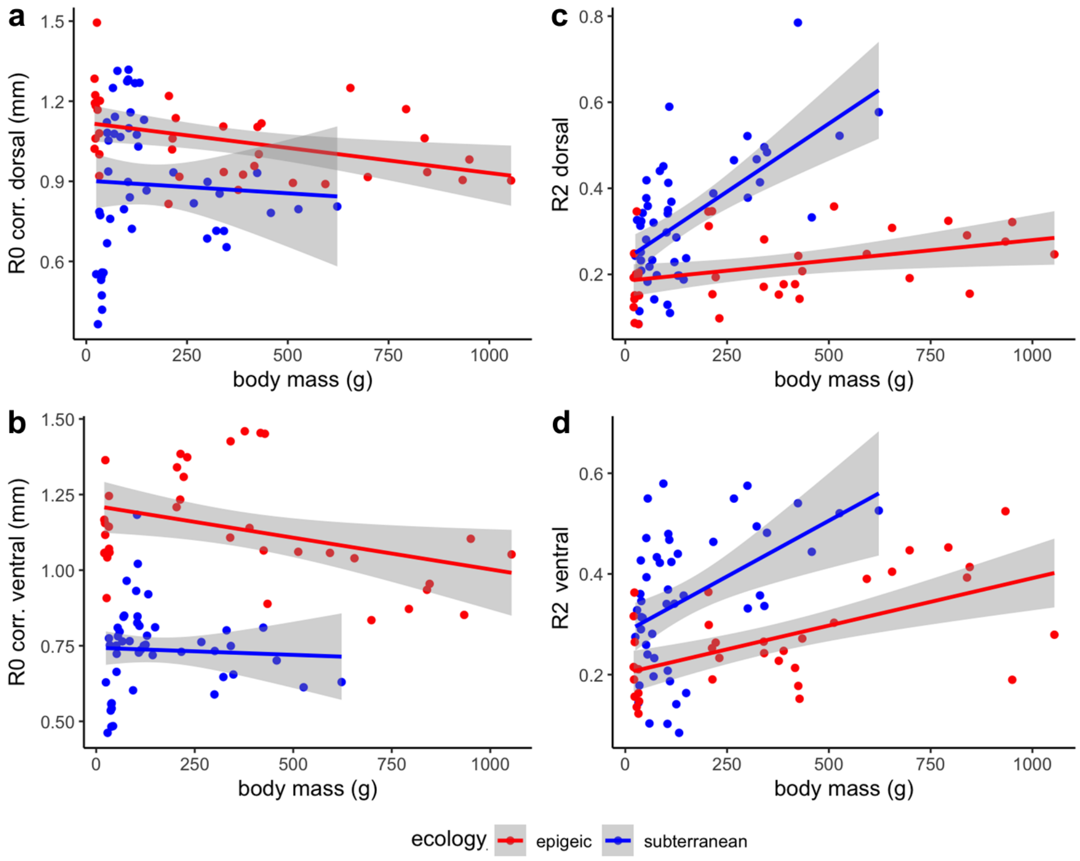


**Supplementary Figure S4**: Effect of body mass on parameters R0 (a – dorsal; b -ventral) and R2 (c – dorsal; d - ventral) in epigeic and subterranean rodents. R0 was corrected for the offset value.

**Part III: Uniaxial tensile testing of rodent skin**

*Skin samples and preparation*

We used uniaxial tensile tests to determine characteristics of small mammal skin when put under horizontal strain. We studied six rodent species, including epigeic and subterranean representatives. For each species, sample sizes, ecology, mean skin thickness, mean max. stress tolerated and mean strain at maximum stress are listed in Suppl. Table S3. Samples were largely derived from frozen specimens (stored at -20 °C) from the research collection of the Department of General Zoology at the University of Duisburg-Essen. Tissue was thawed in sealed falcon tubes prior to testing. Skin flaps of laboratory mice and rats were donated by the animal facilities of the University Clinic of Essen and a portion of NMR samples derived from the Leibniz Institute for Zoo and Wildlife Research in Berlin. The few freshly prepared samples (exclusively laboratory mice, C57BL/6) were tested immediately after animals were sacrificed for other research projects (see Suppl. Table S3). We pooled data from fresh and previously frozen skin samples, since previous research has indicated that maximum stress endured by rodent skin during uniaxial tensile tests is not notably altered by freezing (Foutz et al., 1992).

For testing, frozen specimens were thawed and skin was excised from the back and the ventrum, respectively. The subcutis, including the panniculus carnosus, was carefully removed. The animals' pelage, if present, was left intact to avoid damaging the skin. Later, samples were cut into rectangular shape to fit the clamping jaws of the testing machine (details on the clamping jaws are included in Suppl. Figure S5). Before mounting, the skin thickness was measured at the medial anterior portion of the skin slap utilizing calipers (Format, series: 0600 169). This measurement served as an approximation of the overall thickness of the skin. Sample width varied between 20 mm and 30 mm, with a linear mixed effects model demonstrating that this variable had no significant effect on maximum stress when controlling for species and sample thickness (model structure: max. stress ~ width + thickness + (1|species); $\beta = 0.154$, SE = 0.112, $p = 0.18$). Metric data for all skin samples are listed in Suppl. Table S4 .

A total of 58 skin samples were successfully tested, meaning that no slippage due to incorrect mounting occurred, including 30 dorsal and 28 ventral samples. Of these, 14 samples were collected from epigeic and 44 from subterranean species. This imbalance was due to the availability of the different species on site.

**Supplementary Table S3**: Samples used for uniaxial tensile tests. SD is provided in brackets. *Only one of these samples was successfully tested because of issues during mounting.

| Species | Ecology | Region | *N* | Mean thickness (mm) | Mean max. stress (MPa) | Mean strain at max. stress (%) |
|---|---|---|---|---|---|---|
| *Fukomys anselli* | subterranean | Dorsum | 8 | 0.50 (0.19) | 3.91 (2.53) | 40 (8.25) |
| | | Ventrum | 7 | 0.28 (0.12) | 4.51 (2.12) | 39.8 (6.93) |
| *Fukomys mechowii* | subterranean | Dorsum | 6 | 0.76 (0.27) | 6.18 (2.82) | 83 (28.80) |
| | | Ventrum | 7 | 0.40 (0.07) | 4.96 (1.75) | 63 (18.77) |
| *Heterocephalus glaber* | subterranean | Dorsum | 5 | 0.26 (0.04) | 9.28 (2.73) | 37.4 (7.54) |
| | | Ventrum | 3 | 0.20 (0.03) | 8.6 (1.15) | 40.3 (5.86) |
| *Mus musculus* (C57BL/6) | epigeic | Dorsum | 3 | 0.46 (0.17) | 2.23 (0.06) | 41 (6.0) |
| | | Ventrum | 4 | 0.31 (0.04) | 2.58 (0.93) | 36 (5.69) |
| *Mus musculus* (NMRI-nu) | epigeic | Dorsum | 2 | 0.47 (0.03) | 5.1 (1.41) | 34 (3.0) |
| | | Ventrum | 2* | 0.26 (0.01) | 2.4 | 34 |
| *Rattus norvegicus* (LEW/Crl) | epigeic | Dorsum | 2 | 1.04 (0.01) | 9.0 (0.71) | 115.5 (20.51) |
| | | Ventrum | 2 | 0.41 (0.00) | 9.0 (1.13) | 65 (16.98) |
| *Spalacopus cyanus* | subterranean | Dorsum | 4 | 0.54 (0.28) | 5.22 (2.7) | 44.8 (13.0) |
| | | Ventrum | 4 | 0.44 (0.08) | 3.88 (3.4) | 45.2 (14.5) |

*Uniaxial tensile tests*

Tensile tests were performed with a universal testing machine (Zwick/Roell, Z050, maximum stress = 50 kN). Applied force was monitored by aid of a class 0.5 force transducer (A.S.T. Angewandte Systemtechnik GmbH, type = KAP-S, maximum stress = 2 kN). Skin samples were fixed in an anteroposterior orientation between two customized clamping jaws (see Suppl. Figure S5) with a free clamping length of 16 mm. Subsequently, they were stretched at a constant testing speed of 20 mm/min. Testing was automatically terminated when a sudden drop in force (1 % drop in force) or increase in strain (10 % leap in elongation) was detected, indicating perforation of the sample.

During the experiments, the software of the testing machine (testXpert III, Zwick Roell) would generate a stress-strain diagram (loading curve). Loading curves of mammalian skin (Suppl. Figure S6) exhibit three sections (Silver et al. 2003): a low modulus region in which the tissue offers little resistance to strain, followed by a linear region and a terminal region representing eventual rupture of the tissue. The yield point marks the transition from a reversible deformation (elastic stretching; low modulus region) to a permanent deformation (overstretching of tissue resulting in structural damage; linear region). If the stress is further increased, the tissue will eventually give way (maximum stress/strain) and rupture. Maximum applied stress and strain values were automatically exported by the software.

Given the in parts very small sample sizes per species, we refrained from performing statistical comparisons.

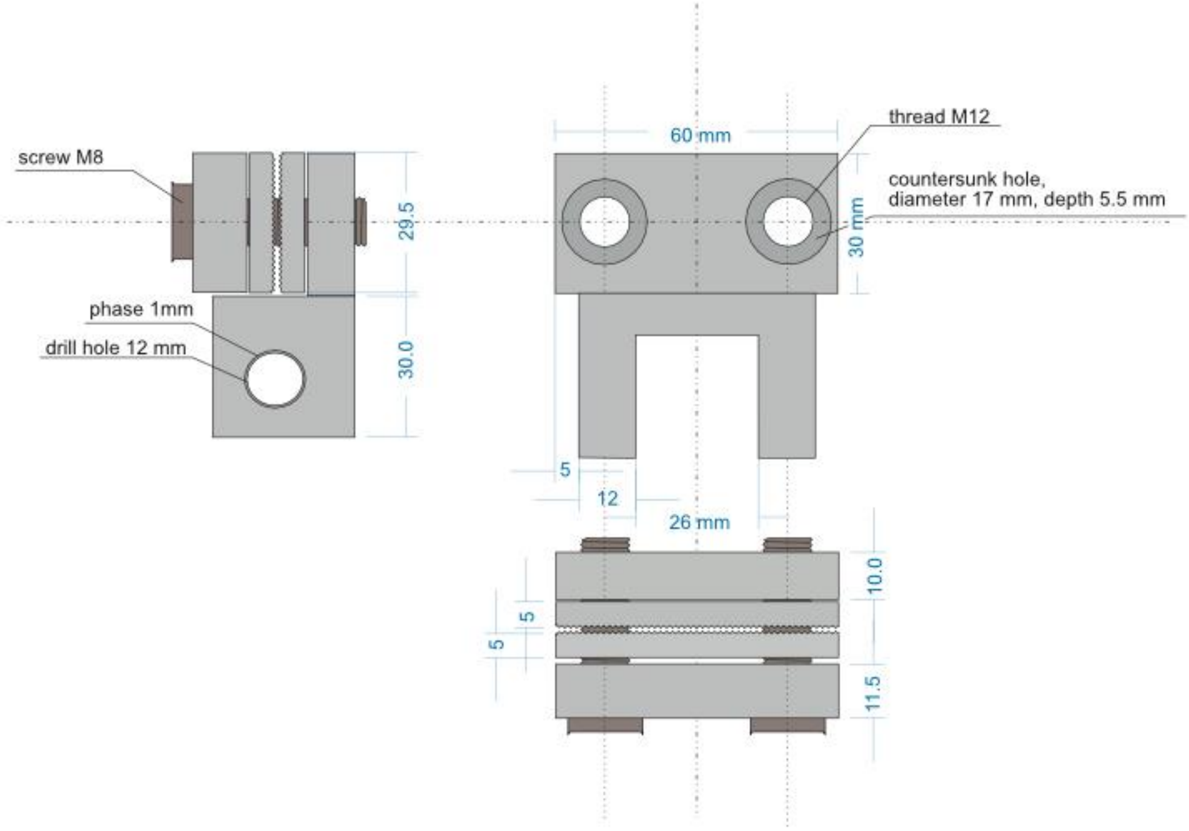


**Supplementary Figure S5**: Design and measurements of customized clamping jaws used for uniaxial tensile tests. Mechanical stress was measured in MegaPascal (MPa), which corresponds to the quotient of the applied force (given in Newton) and the cross-sectional area of the skin sample in square millimeters (1 MPa = 1 million Pa; Pa = 1 N/mm$^2$). Strain was quantified as the relative elongation of the sample (%) compared to its original length.

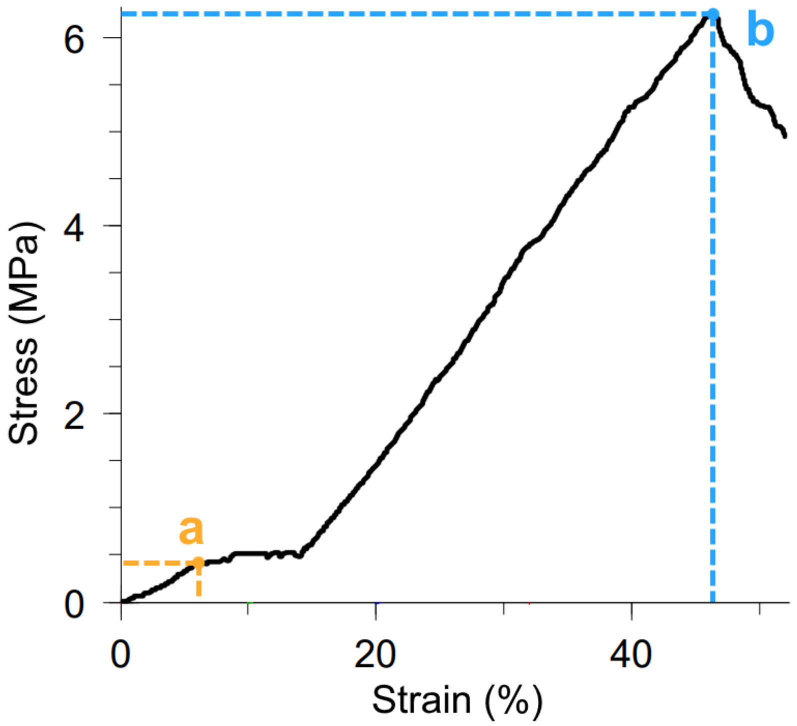


**Supplementary Figure S6**: Exemplary loading curve from a dorsal skin sample of an Ansell's mole-rat (*Fukomys anselli*) showing the triphasic pattern observed in small mammal skin (data collected by the authors). An initial continuous increase is terminated at the yield point (**a**), which marks the end of elastic elongation. Up to the yield point, stress-induced deformation is reversible. Following the yield point, plastic deformation sets in until maximum stress is reached (**b**), after which the sample ruptures.

**Supplementary Table S4**: Individual measurements and biomechanical properties of rodent skin samples subject to uniaxial tensile tests.

| species | region | thickness (mm) | max. stress (MPa) | max. strain (%) | sample width (mm) | Ecology | preservation |
|---|---|---|---|---|---|---|---|
| *Fukomys anselli* | dorsal | 0.83 | 1.6 | 42 | 30 | subterranean | frozen, -20° C |
| *Fukomys anselli* | dorsal | 0.43 | 1.6 | 30 | 23 | subterranean | frozen, -20° C |
| *Fukomys anselli* | dorsal | 0.4 | 2.8 | 28 | 23 | subterranean | frozen, -20° C |
| *Fukomys anselli* | dorsal | 0.67 | 2.1 | 51 | 23 | subterranean | frozen, -20° C |
| *Fukomys anselli* | dorsal | 0.64 | 2 | 49 | 30 | subterranean | frozen, -20° C |
| *Fukomys anselli* | dorsal | 0.42 | 5.6 | 37 | 30 | subterranean | frozen, -20° C |
| *Fukomys anselli* | dorsal | 0.26 | 9.5 | 44 | 30 | subterranean | frozen, -20° C |
| *Fukomys anselli* | dorsal | 0.34 | 3.72 | 39 | 23 | subterranean | frozen, -20° C |
| *Fukomys mechowii* | dorsal | 0.57 | 9.9 | 91 | 30 | subterranean | frozen, -20° C |
| *Fukomys mechowii* | dorsal | 0.76 | 7.4 | 115 | 30 | subterranean | frozen, -20° C |
| *Fukomys mechowii* | dorsal | 1.2 | 2.3 | 89 | 30 | subterranean | frozen, -20° C |
| *Fukomys mechowii* | dorsal | 0.9 | 3.5 | 106 | 30 | subterranean | frozen, -20° C |
| *Fukomys mechowii* | dorsal | 0.47 | 7.7 | 56 | 30 | subterranean | frozen, -20° C |
| *Fukomys mechowii* | dorsal | 0.63 | 6.3 | 41 | 30 | subterranean | frozen, -20° C |
| *Heterocephalus glaber* | dorsal | 0.3 | 13.4 | 34 | 30 | subterranean | frozen, -20° C |
| *Heterocephalus glaber* | dorsal | 0.2 | 8.3 | 41 | 30 | subterranean | frozen, -20° C |
| *Heterocephalus glaber* | dorsal | 0.3 | 9.9 | 48 | 30 | subterranean | frozen, -20° C |
| *Heterocephalus glaber* | dorsal | 0.24 | 8.9 | 36 | 30 | subterranean | frozen, -20° C |
| *Heterocephalus glaber* | dorsal | 0.24 | 5.9 | 28 | 30 | subterranean | frozen, -20° C |
| *Mus musculus* (C57BL/6) | dorsal | 0.65 | 2.2 | 41 | 30 | epigeic | fresh |
| *Mus musculus* (C57BL/6) | dorsal | 0.38 | 2.2 | 47 | 30 | epigeic | fresh |
| *Mus musculus* (C57BL/6) | dorsal | 0.35 | 2.3 | 35 | 30 | epigeic | fresh |
| *Rattus norvegicus* | dorsal | 1.42 | 8.5 | 130 | 30 | epigeic | frozen, -20° C |
| *Rattus norvegicus* | dorsal | 1.42 | 9.5 | 101 | 20 | epigeic | frozen, -20° C |
| *Spalacopus cyanus* | dorsal | 0.78 | 1.7 | 38 | 23 | subterranean | frozen, -20° C |
| *Spalacopus cyanus* | dorsal | 0.34 | 7 | 36 | 23 | subterranean | frozen, -20° C |
| *Spalacopus cyanus* | dorsal | 0.25 | 7.1 | 33 | 23 | subterranean | frozen, -20° C |

| | | | | | | | |
|---|---|---|---|---|---|---|---|
| *Spalacopus cyanus* | dorsal | 0.77 | 2.8 | 54 | 23 | subterranean | frozen, -20° C |
| *Mus musculus* (NMRI-nu) | dorsal | 0.49 | 4.1 | 37 | 30 | epigeic | frozen, -20° C |
| *Mus musculus* (NMRI-nu) | dorsal | 0.45 | 6.1 | 31 | 30 | epigeic | frozen, -20° C |
| *Fukomys anselli* | ventral | 0.35 | 1.7 | 28 | 23 | subterranean | frozen, -20° C |
| *Fukomys anselli* | ventral | 0.42 | 1.9 | 37 | 23 | subterranean | frozen, -20° C |
| *Fukomys anselli* | ventral | 0.43 | 6.3 | 46 | 30 | subterranean | frozen, -20° C |
| *Fukomys anselli* | ventral | 0.28 | 4.2 | 35 | 30 | subterranean | frozen, -20° C |
| *Fukomys anselli* | ventral | 0.21 | 4.6 | 43 | 30 | subterranean | frozen, -20° C |
| *Fukomys anselli* | ventral | 0.17 | 7.4 | 48 | 30 | subterranean | frozen, -20° C |
| *Fukomys anselli* | ventral | 0.11 | 5.5 | 41.3 | 23 | subterranean | frozen, -20° C |
| *Fukomys mechowii* | ventral | 0.38 | 7.6 | 54 | 30 | subterranean | frozen, -20° C |
| *Fukomys mechowii* | ventral | 0.4 | 5.7 | 52 | 30 | subterranean | frozen, -20° C |
| *Fukomys mechowii* | ventral | 0.48 | 3.4 | 77 | 30 | subterranean | frozen, -20° C |
| *Fukomys mechowii* | ventral | 0.4 | 4.1 | 67 | 30 | subterranean | frozen, -20° C |
| *Fukomys mechowii* | ventral | 0.43 | 3.2 | 67 | 30 | subterranean | frozen, -20° C |
| *Fukomys mechowii* | ventral | 0.4 | 3.9 | 33 | 30 | subterranean | frozen, -20° C |
| *Fukomys mechowii* | ventral | 0.25 | 6.8 | 91 | 30 | subterranean | frozen, -20° C |
| *Heterocephalus glaber* | ventral | 0.23 | 9.7 | 38 | 30 | subterranean | frozen, -20° C |
| *Heterocephalus glaber* | ventral | 0.2 | 7.4 | 47 | 30 | subterranean | frozen, -20° C |
| *Heterocephalus glaber* | ventral | 0.18 | 8.7 | 36 | 30 | subterranean | frozen, -20° C |
| *Mus musculus* (C57BL/6) | ventral | 0.35 | 1.3 | 42 | 30 | epigeic | fresh |
| *Mus musculus* (C57BL/6) | ventral | 0.28 | 3.1 | 38 | 30 | epigeic | fresh |
| *Mus musculus* (C57BL/6) | ventral | 0.27 | 3.4 | 33 | 30 | epigeic | fresh |
| *Mus musculus* (C57BL/6) | ventral | 0.32 | 2.5 | 29 | 20.5 | epigeic | fresh |
| *Rattus norvegicus* | ventral | 0.89 | 8.2 | 53 | 30 | epigeic | frozen, -20° C |
| *Rattus norvegicus* | ventral | 0.89 | 9.8 | 77 | 30 | epigeic | frozen, -20° C |
| *Spalacopus cyanus* | ventral | 0.48 | 2.1 | 31 | 23 | subterranean | frozen, -20° C |
| *Spalacopus cyanus* | ventral | 0.52 | 3.6 | 58 | 23 | subterranean | frozen, -20° C |
| *Spalacopus cyanus* | ventral | 0.42 | 9.8 | 53 | 23 | subterranean | frozen, -20° C |
| *Spalacopus cyanus* | ventral | 0.34 | 1.6 | 56 | 23 | subterranean | frozen, -20° C |
| *Mus musculus* (NMRI-nu) | ventral | 0.26 | 2.4 | 34 | 30 | epigeic | frozen, -20° C |
